# Influence of feedback transparency on motor imagery neurofeedback performance: the contribution of agency

**DOI:** 10.1101/2024.02.27.582270

**Authors:** Claire Dussard, Léa Pillette, Cassandra Dumas, Emeline Pierrieau, Laurent Hugueville, Brian Lau, Camille Jeunet-Kelway, Nathalie George

**Affiliations:** Sorbonne Université, Institut du Cerveau - Paris Brain Institute - ICM, Inserm, CNRS, APHP, Hôpital de la Pitié Salpêtrière, Paris, France; Université de Rennes, CNRS, IRISA, UMR 6074, 35000 Rennes, France; Université de Bordeaux, CNRS, EPHE, INCIA, UMR5287, 33000 Bordeaux, France; Institut du Cerveau, ICM, Inserm, U1127, CNRS, UMR 7225, Sorbonne Université, CENIR, Centre MEG-EEG, Paris, France

**Keywords:** Motor Imagery, Neurofeedback, Feedback transparency, Sense of Agency, Beta desynchronisation, EEG

## Abstract

**Objective:** Neurofeedback (NF) is a cognitive training procedure based on real-time feedback (FB) of a participant’s brain activity that they must learn to self-regulate. A classical visual FB delivered in a NF task is a filling gauge reflecting a measure of brain activity. This abstract visual FB is not transparently linked—from the subject’s perspective—to the task performed (e.g., motor imagery). This may decrease the sense of agency, that is, the participants’ reported control over FB. Here, we assessed the influence of FB transparency on NF performance and the role of agency in this relationship.

**Approach:** Participants performed a NF task using motor imagery to regulate brain activity measured using electroencephalography. In separate blocks, participants experienced three different conditions designed to vary transparency: FB was presented as either 1) a swinging pendulum, 2) a clenching virtual hand, 3) a clenching virtual hand combined with a motor illusion induced by tendon vibration. We measured self-reported agency and user experience after each NF block.

**Main results:** We found that FB transparency influences NF performance. Transparent visual FB provided by the virtual hand resulted in significantly better NF performance than the abstract FB of the pendulum. Surprisingly, adding a motor illusion to the virtual hand significantly decreased performance relative to the virtual hand alone. When introduced in incremental linear mixed effect models, self-reported agency was significantly associated with NF performance and it captured the variance related to the effect of FB transparency on NF performance.

**Significance:** Our results highlight the relevance of transparent FB in relation to the sense of agency. This is likely an important consideration in designing FB to improve NF performance and learning outcomes.

## 1. Introduction

Neurofeedback (NF) is a cognitive training procedure that consists of providing real-time feedback (FB) on a participant’s brain activity. The aim is to train them to self-regulate specific brain activity patterns related to a given cognitive ability (Sitaram et al., 2017). For instance, in motor imagery-based electroencephalography NF (MI-EEG NF), participants are trained to reduce the power of sensorimotor activity by imagining movements, since movement preparation, execution, and imagination are associated with power reduction of sensorimotor rhythms (Neuper et al., 2006; Dekleva et al., 2024). These rhythms are typically observed between 8 and 30 Hz over the central regions contralateral to the imagined movement, which we refer to broadly as β activity (Brown & Williams, 2005). In this context, NF performance refers to how well a participant achieves β activity reduction. Despite recent progress, NF performance remains highly variable, with up to 38% of NF participants failing to learn how to self-regulate the targeted activity (Alkoby et al., 2018). Improving FB transparency may be one way to close this NF performance gap.

The type of FB provided during NF training is crucial to promote learning (Lotte et al., 2013). Providing FB improves NF performance compared to MI without FB (Zich et al., 2015), although FB design varies considerably across studies (Roc et al., 2020). While FB is recommended to be clear and meaningful it often takes the form of an abstract stimulus unrelated to the imagined movement, such as a bar that fills based on β activity regulation or MI detection by a classifier. This lacks task-FB transparency, obscuring the causal link between the MI performed and the FB (Beursken, 2012). In contrast, using transparent, task-relevant FB might facilitate MI (Alimardani et al., 2018). It may foster the sense of agency, which is the sense of control over actions such as FB movements in NF (Vlek et al., 2014). Agency is rooted in the consistency between predicted and actual sensory outcomes (David et al., 2008). This may be key to NF learning by fostering task engagement (Jeunet et al., 2016), which is essential to reinforcement learning (Strehl, 2014). Thus, high task-FB transparency may improve NF performance by increasing the sense of agency.

### FB transparency and NF performance

To our knowledge, only one NF study examined the effect of FB transparency on NF performance, comparing a virtual hand FB to an abstract bar FB in a small-sample between-subject study (Ono et al, 2013). They found no improvement of NF performance with the virtual hand FB. A few BCI studies also compared those two types of FB, but their results may not transfer directly to NF, because BCI and NF paradigms typically refer to different types of control metrics. BCI performance refers to task classification accuracy, based on EEG features maximising the contrast between two (or more) tasks, such as left and right MI (Pfurtscheller & Neuper, 2001). It does not place direct emphasis on β activity reduction as is the case in NF. Those BCI studies have obtained mixed results. Skola & Liarokapis (2018) and Penaloza et al. (2018) found that virtual hand FB was associated with better BCI performance than abstract FB, but other studies found no improvement (Neuper et al., 2009; Vourvopoulos et al., 2016).

Multimodal FB is another approach that can be used to increase transparency. Robotic hand orthoses can provide FB by passively moving the participant’s hand. They have shown encouraging results mostly on BCI performance, as compared to simple bar FB (Vukelić & Gharabaghi, 2015; Darvishi et al., 2017; see also Ono et al., 2018). However, they induce real movements, which can modulate the targeted β activity, making their use in a NF context challenging. An alternative approach is to use tendon vibrations to mimic the perception of the imagined movement by generating motor illusions (Roll & Vedel, 1982). Such vibrations were associated with higher BCI performance and a longer-lasting 8-13Hz ERD when combined with a virtual hand FB (Barsotti et al., 2018).

Overall, previous research has provided evidence that FB transparency may improve NF performance, but the underlying mechanisms remain unclear. We hypothesize that this may be related to the influence of FB transparency on agency.

### Agency and FB transparency in EEG NF

Agency is the sense of control over one’s actions. It can extend to external objects in a disembodied way through motor actions like keypresses (Caspar et al., 2015; Zopf et al., 2018). It has recently emerged as a key concept in human-computer interaction, especially in neurotechnology (Schönau et al, 2021) and brain-computer interfaces (Kostick et al, 2021; Nierula et al, 2021; Venot et al, 2024). Yet, research on the sense of agency in mental tasks (not related to overt motor actions) is limited. In BCI and NF studies, the FB is the external object controlled by the participant. No NF study has investigated the effect of FB transparency on agency. Based on the comparator model of agency (Frith et al, 2000), one could predict that FB transparency, in the form of the task-FB matching would increase the sense of agency. In a related line, a few BCI studies reported that various aspects of FB design influenced agency. Evans et al. (2015) manipulated the congruency between the movement direction of a visual FB formed by an abstract cursor and the MI side. They reported that congruency drove sense of agency in this BCI paradigm. This result was recently replicated and extended to somatosensory FB in an invasive intracortical BCI case study (Serino et al., 2022). These authors used virtual hands as visual FB and neuromuscular stimulation as somatosensory FB. By manipulating congruency for each FB modality, they showed that agency was influenced by both visual and somatosensory FB cues with a primacy of the latter (Serino et al., 2022). Finally, a BCI study compared agency levels between a MI-based task and steady-state visual evoked potential (SSVEP) based task, keeping FB identical in both tasks, in the form of a virtual hand. They found higher agency over FB in the MI task, corresponding to the more transparent condition (Nierula et al, 2021).

Here, we sought to test the link between transparency, NF performance, and agency in a MI-EEG NF protocol. We created a NF protocol with three different FB conditions designed to provide different levels of transparency. These FB took the form of a swinging pendulum, a clenching virtual hand, and a clenching virtual hand combined with motor illusion-inducing vibrations. The amplitude of pendulum and hand movements as well as the delivery of vibrotactile stimuli were based on the amount of β activity reduction measured on the central electrode contralateral to the imagined hand movement. Recordings of MI alone and passive conditions of FB stimulus presentation were included to disentangle the effect of MI from movement observation / vibrotactile stimulation *per se.* We tested the following hypotheses: Transparency improves NF performance (H1); transparency increases judgment of agency (H2); agency captures a large part of the variance related to the effect of transparency on NF performance (H3).

## 2. Material and methods

### 2.1 Participants

Twenty-five healthy human participants were included in the study (12 women; age: 27.8 ± 6.9 years [mean ± SD]; 24 right-handed and 1 ambidextrous, according to self-report). All participants had normal or corrected-to-normal vision, reported no history of psychiatric or neurological pathology and no consumption of alcohol or psychoactive drugs in the week prior to the experiment. All participants provided written informed consent to participate and received 80 € for participating. The study was approved by the CPP Ile de France VI local ethics committee and was carried out in accordance with the Declaration of Helsinki. Two participants were excluded from the analyses (one for non-compliance with the instructions and one for technical difficulties). Thus, we analysed the data of 23 participants (11 women, 12 men).

Sample size was determined with reference to a study with comparable methods (Ziadeh et al., 2021) in which 22 participants were tested. Our study with 23 participants resulted in a sensitivity to detect effect sizes of at least 0.04 to 0.10 in η^2^, with type I error rate alpha = 0.05 and statistical power of .80, as computed using G*power (version 3.1.9.7) for a 3-conditions within-subject repeated-measures design, with correlation among repeated measures between 0.4 and 0.7, and non-sphericity correction between 0.7 and 1.

### 2.2 Description of FB stimuli

We used three types of FB (Figure 1A). The first two FB were visual and represented either a pendulum (PENDUL condition) or two virtual hands (HAND condition). The third FB was multimodal, featuring both the virtual hand visual stimuli and tactile vibrations eliciting motor illusion of right-hand closing (HAND+VIB condition). All these FB conditions were driven by the modulation of the targeted β activity (see *2.4 Online EEG signal processing* for details).

**Figure 1.**
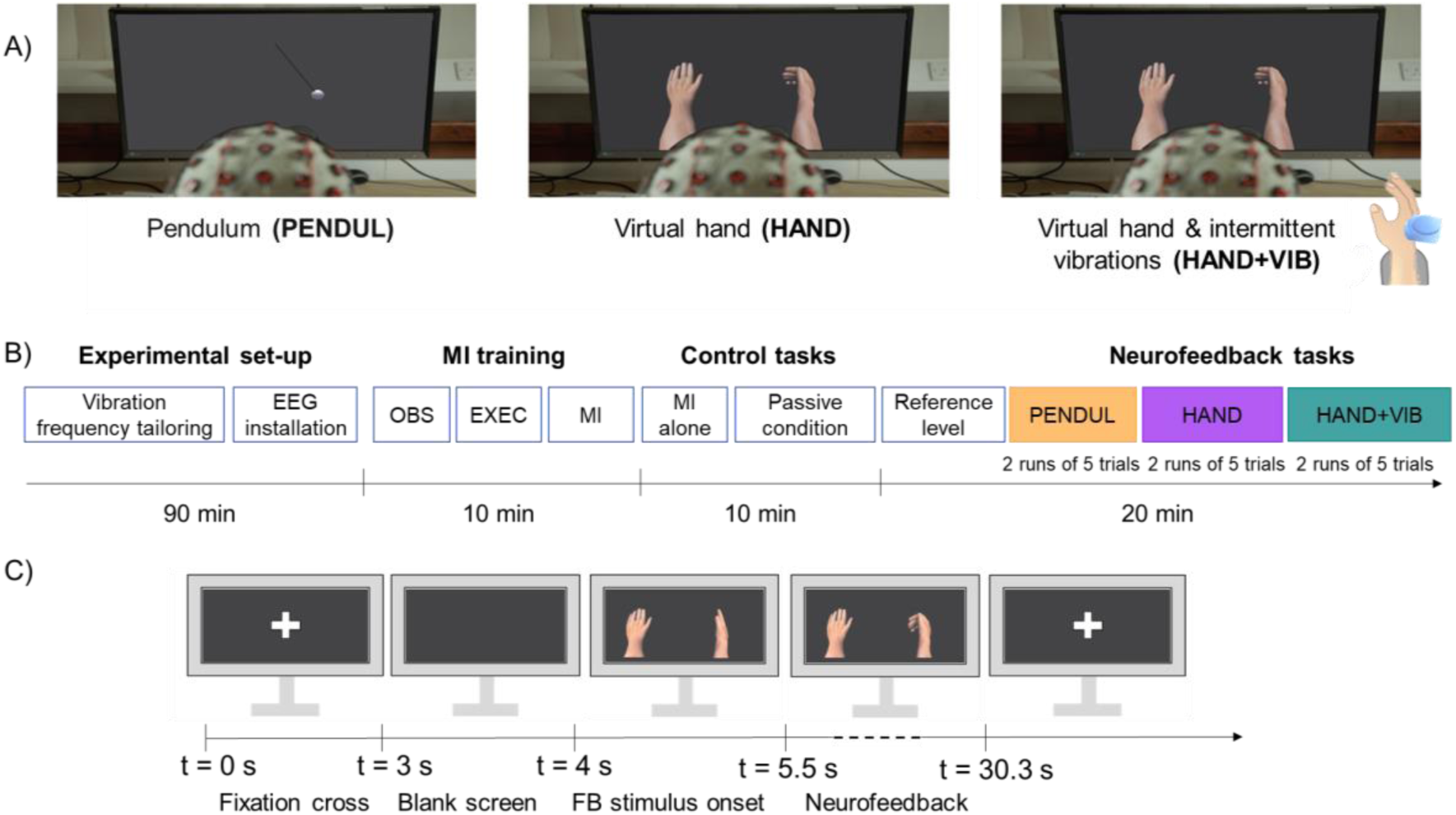
Illustration of the experimental protocol. ***A*. Representation of the different FB conditions**, i.e., pendulum (PENDUL), virtual hand (HAND), and virtual hand with motor illusions FB (HAND+VIB), which we assumed to be of increasing transparency. ***B*. Time course of the experiment.** The experiment began with individual tailoring of the motor illusion vibration frequency followed by EEG cap installation. The participant performed MI training in 3 blocks: observation (OBS), execution (EXEC), and motor imagery (MI). Then, control tasks were performed, including two blocks of MI without any FB (MI alone) and three blocks of passive control tasks (Passive condition). The reference β level for the NF was measured. The NF task included two runs of each FB condition (Pendulum, Virtual hand, Virtual hand with vibrations), performed in counterbalanced order across participants. ***C.* Time course of an example trial in the HAND condition.** The trial started with a fixation cross, followed by a black screen and the onset of the FB static picture. FB stimulus movement started after 1.5 s and lasted for 24.8 s. At the end of the trial, the fixation cross was displayed again until the next trial began. See main text for further details.

The visual FB stimuli were programmed using Unity C# ®. The virtual right hand performed opening and closing movements and the pendulum swayed from the centre to the right side of the screen. Movement amplitude varied as a function of the measured β activity. The pace of the movements was kept constant with one forth and back sway of the pendulum and one opening and closing of the hand every 1.55 s, to help participants imagine the movement in synchronicity with the visual FB (Eaves et al., 2016).

The tactile stimuli were delivered by a VibraSens VB115® (TechnoConcept, Manosque, France) attached using a gauze strip to the back of the participant’s right hand, over the extensor tendons.

### 2.3 Experimental protocol

The experiment consisted of a single NF session with three blocks of different FB modalities corresponding to three different levels of transparency. The experimental protocol is described in Figure 1B. First, the vibrator was attached to the participant’s right hand, which rested on an armrest, hidden from sight by a table. Vibration frequency was individually tailored between 65 and 90 Hz to elicit the motor illusion of a closing hand (mean across participants = 69 Hz ± 9 Hz) (Taylor et al., 2017). The vibrator was kept in place throughout the experiment so that recording conditions were comparable.

The EEG recording system was then installed. Signal quality was visually inspected, and we aimed for electrode impedances below 10kΩ (median across electrodes = 11 ± 10 kΩ). The participants were seated 80 cm away from the screen in a dimly lit electrically shielded room and instructed to avoid moving during the EEG recordings.

Participants were first familiarised with the MI task through a 30s-video which demonstrated rhythmic opening and closing movements of a real hand. Training consisted of three 30 s phases. First, the participants observed the movement to understand its nature and rhythm (OBS in Figure 1B). Then, they executed the movement with their right hand at the same rhythm (opening and closing cycles of 1.55 s) as the video (EXEC in Figure 1B). Finally, the participants imagined the movement while watching the video (MI in Figure 1B). This training was repeated until participants felt comfortable with the task (one repetition for most participants). Kinaesthetic imagery was encouraged by emphasising the importance of feeling the sensations associated with the imagined movements as it has been associated with better MI performance (Neuper et al., 2005).

The participants performed two 30 s sequences of MI without any FB, with only a fixation cross displayed at the centre of the screen (MI alone condition in Figure 1B). Furthermore, as movement observation and vibrotactile stimulations can influence *β* activity, we included three 30 s sequences of passive control tasks. During these control tasks, participants remained at rest while being presented with random pendulum, random virtual hand movements, or random virtual hand movements with intermittent motor illusion vibrations (one 30 s block per condition). For technical reasons, the MI alone condition was performed by 20 out of the 23 participants and the passive control conditions by 21 out of the 23 participants.

Before starting the NF training, we recorded baseline EEG activity to determine a *reference β* power level. Participants were instructed to remain still and relaxed while fixating on a central cross on the screen, during two runs of 30 s each. This enabled us to set the individual *reference β* level required to compute FB for the rest of the experiment (see below). Then, NF training started. Participants completed two consecutive runs of each FB condition. The presentation order of the FB conditions followed a Latin square design to ensure counterbalancing across participants. Each run consisted of 5 trials of 30 s each. A trial unfolded as follows (Figure 1C):

At t = 0s, a fixation cross was displayed for 3 s, followed by a 1s black screen. At t = 4s, motionless pendulum or hands were displayed for 1.5 s while participants started performing MI. From t = 5.5 to t=30.3 s (duration of 24.8 s), visual FB was continuously provided while participants performed MI. In HAND+VIB, additional vibrotactile FB could be provided for ∼2 s at t=10.4 s, 16.7 s, 22.7 s, 28.9 s. We designed it as an intermittent FB to avoid sensory habituation leading to opposite direction movement illusions (Taylor et al., 2017). Every trial featured 16 visual FB movement cycles of 1.55 s. At the end of the trial, the fixation cross was displayed again. The next trial began after a random inter-trial time interval of 3 to 4.5 s.

Each run of 5 trials was followed by a 1-minute break where participants filled out questionnaires (see below).

### 2.4 EEG data acquisition and signal processing

Electroencephalographic (EEG) signal was recorded using a 32 active electrode cap (ActiCAP snap, Brain Products GmbH) with actiCHamp Plus system (Brain Products GmbH), with electrodes placed according to the extended international 10-20 system (Fp1, Fp2, F7, F3, Fz, F4, F8, FT9, FC5, FC1, FC2, FC6, FT10, T7, C3, Cz, C4, T8, TP9, CP5, CP1, CP2, CP6, TP10, P7, P3, Pz, P4, P8, O1, O2, Oz). EEG data were recorded at 1kHz with a band-pass filter of DC-280Hz by BrainVision Recorder. The signal was transmitted to the OpenViBE 2.2.0 acquisition server (Renard et al., 2010) for online processing and FB computation using the Brain Products Brainamp Series driver. The signal was recorded with reference to Fz and the ground electrode was placed on Fpz.

Electrooculographic (EOG) activity was recorded using two bipolar disposable electrodes, placed above and below the right eye for vertical eye movements, and on the right and left outer canthi for horizontal eye movements. Electromyographic (EMG) was recorded from the right hand using two disposable electrodes placed on the inner side of the right wrist, where small hand movements produced detectable contractions. Electrocardiographic (ECG) activity was recorded using two disposable electrodes placed on the right clavicle and the left lower abdomen. A disposable electrode placed on the left shoulder of the participant was used as the ground for the eight EOG / EMG / ECG electrodes.

#### Online EEG signal processing for NF computation

The EEG processing pipeline used in OpenViBE 2.2.0 to compute the β activity that the participants aimed to regulate through NF is described below. A Laplacian spatial filter was computed over the C3 electrode by subtracting the neighbouring signals arising from CP5, CP1, FC1 and FC5 (Blankertz et al., 2007). The resulting EEG signal was band-pass filtered between 8 and 30 Hz using an 8^th^ order, 1-way, infinite impulse response (IIR), Butterworth filter. The signal was then epoched in 1s time windows with 0.75 s overlap. For each epoch, the EEG signal was squared and averaged over time, to obtain the epoch β band power value. From t = 4 s in each trial, these values were streamed using the Lab Streaming Layer (LSL) communication protocol (Kothe et al, 2024) to a Unity application to provide FB to the participant (see Figure 1B). We averaged *online β* band power values so that each FB cycle amplitude was determined by the average of the 4 consecutive epochs just before the cycle onset as compared to the participant’s *reference β* power (see below). To reduce overlap between means, two epoch values were dropped between each FB cycle (see Supplementary Figure 1A). We used the same pipeline offline to compute *offline β* power values in the control conditions.

#### Reference β power

*Reference β* power was computed using data from a 2 x 30 s baseline using the same method as described above for online EEG signal processing. The median of the obtained β power values was taken as the *reference β* power, with manual adjustment if needed to allow the participant to receive FB during the task (see Supplementary Table 1).

#### Visual FB

The greater the detected reduction in *online β* power relative to the *reference β* power, the larger the FB movement was. Thresholds for FB movement were set at a 10% and 55% reduction in *online β* power compared to *reference β* power based on previous reports (Crone et al., 1998; Kühn et al., 2006) and confirmed empirically during pretests. Less than 10% *online β* power reduction led to no FB movement on screen. More than 55% *online β* power reduction led to a full amplitude FB movement on screen. Between these lower and upper limits, amplitude was proportional to *online β* power reduction. The pace of the hand/pendulum FB movements was kept constant (with a cycle duration of 1.55s) to help participants imagine at the trained pace, in synchronicity with the visual FB.

#### Tactile FB

Tactile FB was triggered using a Python script and an Arduino Uno ® microcontroller board. Vibrations were triggered a maximum of four times per trial on the 4^th^, 8^th^, 12^th^ and 16^th^ cycles of the visual FB, corresponding to every ∼6.2 s during the trial. This FB was intermittent to avoid sensory habituation and opposite direction illusions (Taylor et al., 2017). The decision to trigger the vibration was based on the values collected during the preceding 3 FB cycles. Thus, tactile FB was based on a longer time window than visual FB. For each vibration, we considered the 15 β band power values (i.e. 1 s epochs) collected during the preceding 3 FB cycles (leaving out 1 epoch i.e. 1s of signal just before vibration to accommodate triggering delay). We computed the weighted mean of these values, giving less weight to the values furthest in time from the vibrations to improve visuo-tactile FB coherence (see Supplementary Figure 1B). We compared this weighted mean to the *β reference* level. The threshold was set to a reduction of 30% relative to the *β reference* level (corresponding approximately to the mean between the 10% and 55% thresholds used for visual FB). Less than 30% reduction led to no vibration while more than 30% reduction triggered a motor illusion vibration, which lasted for ∼2 s.

#### Offline signal processing and analysis

##### NF performance

First, at the behavioural level, for each FB condition, we examined the number of trials with visual FB (aka. FB movement) and the total number of FB movements > 5% maximum amplitude across trials. We also computed the mean amplitude of visual FB movements including all trials, that is, including trials with and without any FB movement. For the HAND+VIB condition, we computed the total number of vibrations delivered and the number of concomitant visual and tactile FB.

Second, we computed NF performance as follows. Every trial featured 16 FB movement cycles based on 16 values of *online β* power compared to the participant’s *reference β* power, as detailed above. Thus, we divided these *online β* power values by the *reference β* power and computed the median of the 80 ratios (16 values per trial x 5 trials per run) for each FB condition and each run, for every participant. We log-transformed the result and took the opposite so that positive NF performance reflected a reduction of *online β* power relative to the *reference β* power. We used the same method to compute the equivalent of NF performance (so-called “offline performance” in the remaining text) for the control conditions (passive and MI alone tasks).

##### Time-frequency analysis of whole-scalp EEG data

We performed offline analysis of the event-related desynchronisation and synchronisation (ERD/ERS) across the frequency spectrum and the whole scalp to validate our experimental protocol. Raw EEG data in BrainVision Recorder format were analysed using the MNE 0.23.0 package (Gramfort et al., 2013) under Python 3.9 environment. A 0.1 Hz high-pass and a 90 Hz low-pass IIR Butterworth filters of 4^th^ order were applied, along with two zero-phase notch filters with stop bands centered at 50 and 100 Hz. Each participant’s signal was epoched into NF trials, from 5 s before to 31 s after the visual FB onset (time 0 being the onset of the visual FB display). We reviewed the trials and removed those with muscle artefacts. On average, 8 out of 10 trials remained in each FB condition, for each participant. Four electrodes located around the maxillary regions (TP9, FT9, TP10, and FT10) presented frequent muscle artefacts and were removed from the analysis. Independent component analysis (ICA) was used to remove ocular artefacts. For this, we applied an additional high-pass filter with 1-Hz cut-off to the raw data and computed ICA on 1 Hz high-pass filtered epochs using fastICA (Hyvärinen & Oja, 2000). We identified two to three components with blink or saccade signatures for each participant. The resulting ICA unmixing matrix was applied to the 0.1 Hz high-pass filtered epochs, subtracting the identified ocular independent components. The data were then average referenced and down-sampled to 250 Hz. We used a Morlet wavelet transform to compute the EEG signal power between 3 and 84 Hz, with 1 Hz frequency bins. The number of wavelet cycles increased linearly with frequency to maintain a constant time/frequency resolution. The resulting time-frequency data were averaged across trials for each participant and FB condition. Finally, power values were normalised relative to baseline using a log-ratio of power at each time point relative to the mean power over 2 s of fixation cross (from -3 to -1 s) before the onset of the FB visual stimuli, for each frequency bin (see Supplementary Figure 4 for a graphical overview of the whole processing pipeline).

The obtained ERD/ERS data were averaged across time, excluding the first and last second of the FB movement time period in each FB condition. Topographical maps of the ERD/ERS data were computed by averaging these data in the targeted 8-30 Hz band, in each FB condition, across subjects. We also computed the standardised effect size of ERD/ERS for each FB condition in the form of Cohen’s d, by dividing the mean ERD/ERS values by the standard deviation across the 23 participants, for each frequency from 3 to 84 Hz and on each of the 28 electrodes.

### 2.5 Questionnaires

After every run of 5 trials for each condition, the participants were asked to report their experience by scoring the following statements inspired from Fribourg et al. (2018) and Nierula et al. (2021) on 11-point Likert scales (0 to 10). Each statement assessed an aspect of the participant’s experience as indicated in the square brackets:

- I felt like the virtual hand was mine. (Only for HAND and HAND+VIB) [ownership]

- I felt like I was the one controlling the movements of the hand/pendulum. [agency]

- I felt like the hand/pendulum was moving by itself. [external causal attribution]

- I felt like the task was difficult. [difficulty]

- I felt like I succeeded at the task. [success]

- I felt like the task was enjoyable. [satisfaction]

Difficulty, success and satisfaction scores were combined to create a user_experience composite score using the first component of the principal component analysis (PCA). This PCA-computed score accounted for 73% of the variance of the 3 Likert scales across participants. The original values of ownership, agency and external causal attribution scores were analysed.

In addition, after the two runs of each condition, the participant filled out a questionnaire to evaluate the FB. This questionnaire was adapted from Kübler et al. (2014) and comprised the following statements, scored on 11-point Likert scales:

- I found this FB easy to use. [ease of use]

- I learned to use this FB quickly. [learnability]

- I enjoyed the FB aesthetics. [aesthetics]

- I found this FB reliable. [reliability]

- I found this FB reactive. [reactivity]

These five scores were combined to create a feedback_usability composite score. This PCA-computed score accounted for 70% of the variance of the 5 Likert scales across participants.

### 2.6 Statistical analyses

#### Experimental validation

First, to validate our experimental manipulation, we evaluated the spatio-spectral pattern of ERD/ERS in each FB condition. We assessed the statistical significance of ERD/ERS against zero in each FB condition using one-sample t-tests performed on each frequency and electrode (*mne.stats.permutation_t_test* function of MNE-Python library). To correct for multiple comparisons across both frequency and electrode dimensions, we performed 20,000 permutation-based t-tests. We selected the maximum t value (tmax) across our 2,296 dimensions (28 electrodes x 82 frequencies) on each permutation to build the FWER-corrected statistical distribution of tmax values under the null hypothesis (FWER = family-wise error rate). The observed t-values were compared to this distribution and considered significant if they belonged to the top 1.67% (5% divided by 3, for 3 FB conditions) of the tmax distribution.

#### H1 – Transparency improves NF performance

We analysed the data using R (version 4.0.4) within the RStudio IDE (version 2022.02.0) and JASP software (version 0.14.1) (Love et al., 2019). The code and data for the analyses is available via OSF at: https://osf.io/es8b7/?view_only=16ba71af1ba5419686d87656fbf5dd75

To test our first hypothesis, we analysed NF performance using a 2-way repeated measures Analysis of Variance (ANOVA) with FB_conditions (PENDUL, HAND, and HAND+VIB) and runs (1 and 2) as within-subject factors. When the degree of freedom of the comparisons was greater than 1, we used Greenhouse-Geisser correction for deviations from the sphericity hypothesis. We reported F values with original degrees of freedom, corrected p values, and effect size in the form of general eta-squared (η^2^). Planned comparisons of FB conditions were performed using unilateral Student t-tests in accordance with our hypothesis that increasing transparency would increase performance (HAND > PENDUL and HAND+VIB > HAND), whenever a significant effect of FB conditions or interactions were found. We report t and p values and Cohen’s d effect size for these tests. For the HAND+VIB vs. HAND comparison, we also report the result of bilateral t-test because an effect opposite to the expected one was observed.

We used a linear mixed effect regression (LMER) model to examine the evolution of NF performance across trials for the different FB conditions. In this analysis, all the values of NF performance along time except the first one were considered for each trial under the three FB conditions (n = 15 values per trial). The data were analysed using the *lmer* function of the lme4 R package. Model structure was determined through nested testing (see supplementary section VI for details). We included FB_conditions, Trials, and Time as fixed-effect factors. We included random intercept and random effect of FB conditions across participants. The LMER model was the following, as coded in R:

NFperformance ∼ FB_conditions * Trials + Time + (1 + FB_conditions | participant_id)

In this model, Time was coded as a numerical factor. Trials was initially coded as a categorical factor and then as a numerical factor to test the linear trend of NF performance across Trials under the three FB Conditions. Model parameters were estimated using the Restricted Maximum Likelihood approach. F and P-values were estimated via type III ANOVA with Satterthwaite’s method using the *anova* function of the R package lmerTest.

We performed post-hoc comparisons of NF performance for the different FB conditions using the emmeans package. For the model where Trials was coded as a numerical factor, we tested the linear trend of NF performance across trials in each FB condition using the emtrends package.

#### Additional control analyses

To control for a potential effect of the order of FB conditions, we performed a 1-way ANOVA using the order of FB conditions (1^st^, 2^nd,^ and 3^rd^ FB conditions) as within-subject factor. We took the median value of NF performance across both runs for this analysis.

Moreover, we conducted additional analyses to examine the impact of artefact sources, specifically electrooculographic (EOG) and electromyographic (EMG) activities (Fatourechi et al., 2007) on our results. These analyses are reported in Supplementary Material section IV.

Finally, we performed a 2-way ANOVA to test for the effect of FB stimuli *per se* on NF performance, with FB_conditions and Task (NF versus passive control tasks of FB stimuli) as within-subject factors. For the analysis of the MI alone control task, we computed the mean value of NF/offline performance across the 3 stimulus conditions (PENDUL, HAND, HAND+VIB) for the Passive and NF tasks and we conducted a one-way ANOVA with Task as within-subject factor (NF / Passive / MI alone).

#### H2 – Transparency increases judgment of agency

We analysed agency ratings using a 2-way repeated-measures ANOVA with FB_conditions and runs as within-subject factors. We conducted planned comparisons of FB conditions using unilateral Student t-tests (HAND > PENDUL and HAND+VIB > HAND) when a significant effect of FB conditions or interaction involving this factor was found. External causal attribution, ownership, and the user_experience score derived from the questionnaires were analysed in the same way. We analysed the feedback_usability score, assessed once after both runs in each FB condition, using a 1-way ANOVA with FB_conditions as within-subject factor.

#### H3 – Agency captures a large part of the variance related to the effect of transparency on performance

For each participant, we had six paired values of NF performance and agency, corresponding to the two runs of each FB condition. To assess the relative influence of Agency and FB_conditions on NF performance, we computed incremental LMER models, with i) either FB_conditions (Model 1) or Agency (Model 3), ii) FB_ Conditions *and* Agency (Model 2), iii) FB conditions, Agency *and* interaction between FB_conditions and Agency (Model 4), as fixed effect factors accounting for NF performance. In all these models, FB_conditions was a categorical factor, with HAND as reference level. Agency was a numerical factor and agency score values were standardized by z-scoring across all values. Run was included as an additional, fixed effect to account for training or fatigue effects. We kept the random effect structure constant with FB_conditions and Agency as random effects across subjects in addition to random intercept across subjects. The incremental models were compared using ANOVA (*anova* function of R stats package). We report chi-square tests of goodness of fit between models (χ²) and associated p values.

## 3. Results

### 3.1 Overall group-level performance

In our NF task, FB could be provided in three different forms (PENDUL, HAND, or HAND+VIB).

Twenty participants out of 23 managed to reduce their *online β* power relative to the *reference β* power so that they triggered movements in PENDUL condition, while 21 participants did for HAND condition and 16 participants triggered vibrations for HAND+VIB condition. Fifteen participants managed to trigger FB in all 3 FB conditions. Table 1 summarises the amount of FB received in each condition across participants and individual data are illustrated in **S**upplementary Figure 2.

**Table 1.** Amount of FB received by the participants in all 3 experimental FB conditions (median/mean ± std across the 23 participants).

| FB condition | PENDUL | HAND | HAND+VIB |
| --- | --- | --- | --- |
| Median number of trials with positive visual FB (out of 10) | $8 \pm 3$ | $9 \pm 3$ | $7 \pm 4$ |
| Median number of >5% amplitude FB movements (out of 160) | $58 \pm 53$ | $86 \pm 47$ | $56 \pm 47$ |
| Mean amplitude of visual FB movement (in % of maximum amplitude) | $24.9 \pm 2.5$ | $33.6 \pm 2.7$ | $20.6 \pm 2.2$ |
| Median number of vibrations (out of 40) | — | — | $9 \pm 13$ |
| Median number of concomitant visual and vibratory FB (out of 40) | — | — | $7 \pm 10$ |

In the HAND +VIB condition, vibratory FB could occur every four cycles of the visual FB only. Its computation resulted in some incongruence with the visual FB, that is, vibrations could occur in the absence of simultaneous visual FB. We evaluated the frequency of congruent (aka. concomitant) and incongruent (aka. vibratory FB without virtual hand movement) visuo-tactile FB in the HAND+VIB condition. Over the total number of 40 possible visual and tactile FB cycles, vibrations occurred 32.3% of the time on average in the absence of visual FB movement (incongruent cases). All sixteen participants who triggered positive tactile FB but one experienced this incongruent case of tactile FB without co-occurring visual FB (Table 1 and Supplementary Figure 3).

### 3.2 Does transparency improve NF performance? (H1)

We validated our experimental manipulation by examining the topographical maps of the ERD/ERS data averaged over the targeted β band. Figure 2A presents the grand average of these maps across participants in each FB condition. The participants successfully modulated the targeted activity, with desynchronisation in the β band observed over the left sensorimotor cortex centred on C3 electrode in all FB conditions. This pattern seemed the most pronounced in the HAND condition (see also **S**upplementary Figure 5 for time-frequency representations).

**Figure 2.**
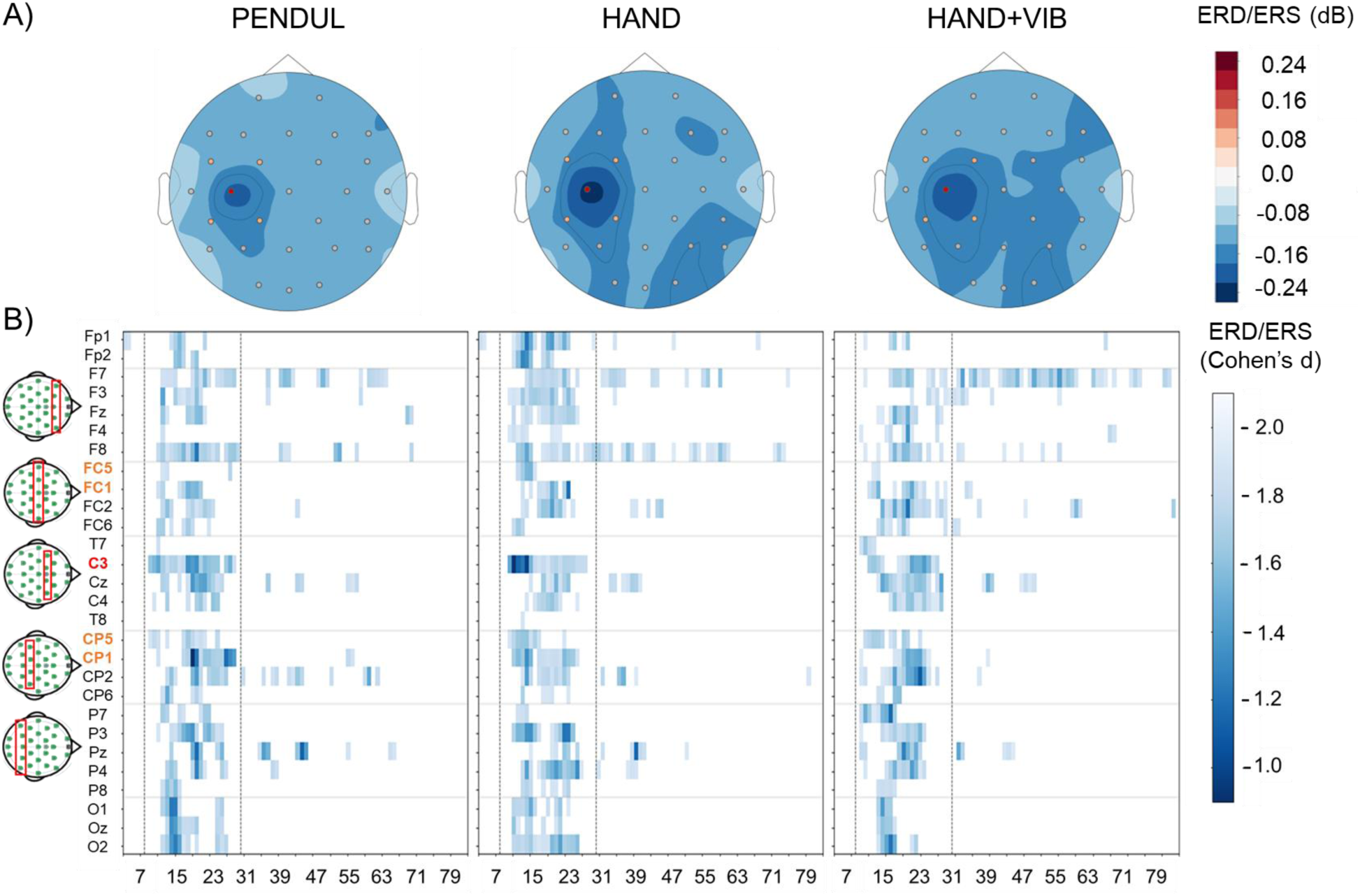
***A.*** Topographical maps of **ERD/ERS in the β band during NF trials for each FB condition.** Grand average in the 8-30Hz band across the 23 participants for PENDUL, HAND, and HAND+VIB. The blue colours represent ERD, red colours represent ERS. ***B.* ERD/ERS effect sizes across frequencies and electrodes, in the 3 FB conditions.** Values are masked by significance, showing only the effect sizes that are significantly different from zero at a corrected p value < 0.05. Significant ERD effect sizes are represented in blue scale on electrode-frequency plots with frequencies in abscissa and electrodes in ordinate. Electrodes are organized in antero-to-posterior transversal montages represented as red rectangles in the left small maps. Laplacian FC1, FC5, CP1, and CP5 electrodes are written in orange, and C3 is in red. The vertical dotted lines delineate the targeted 8-30 Hz frequency range.

We computed the effect size of the ERD/ERS on each frequency and electrode to assess the spatio-spectral selectivity of brain activity modulation across subjects. For the three FB conditions, desynchronisation was focused in the targeted 8-30Hz band (Figure 2B). The mean effect size (Cohen’s d) of ERD on C3 was on average -1.32 for the PENDUL condition, -1.31 for HAND and -1.16 for HAND+VIB condition.

We next tested the influence of FB conditions on NF performance (i.e., *online β* power reduction relative to the *reference β* power). The ANOVA with FB_conditions and runs as within-subject factors revealed a significant main effect of FB_conditions (F(2,44) = 3.91, p = .03, η^2^= .10), as represented in Figure 3. This reflected higher NF performance in HAND (mean, M = 0.07, standard deviation, SD = 0.18) compared to PENDUL (M = -0.01, SD = 0.18) (t(22) = 1.88; p =0.035; d = 0.39). In contrast, NF performance was decreased in HAND+VIB (M = -0.04, SD = 0.21) relative to HAND. Thus, the unilateral t-test of our planned comparison (HAND+VIB > HAND) was not significant, while a bilateral t-test indicated a significant difference in the direction opposite to the expected one (t(22) = -0.64; p = 0.005; d = -0.64). There was a trend towards a decrease in NF performance across runs, but this effect did not reach statistical significance (F(1,22) = 4.08, p = 0.06, η^2^ = .03). There was no significant interaction between FB_conditions and Run (F(2,44) = 0.36, p = 0.7).

**Figure 3.**
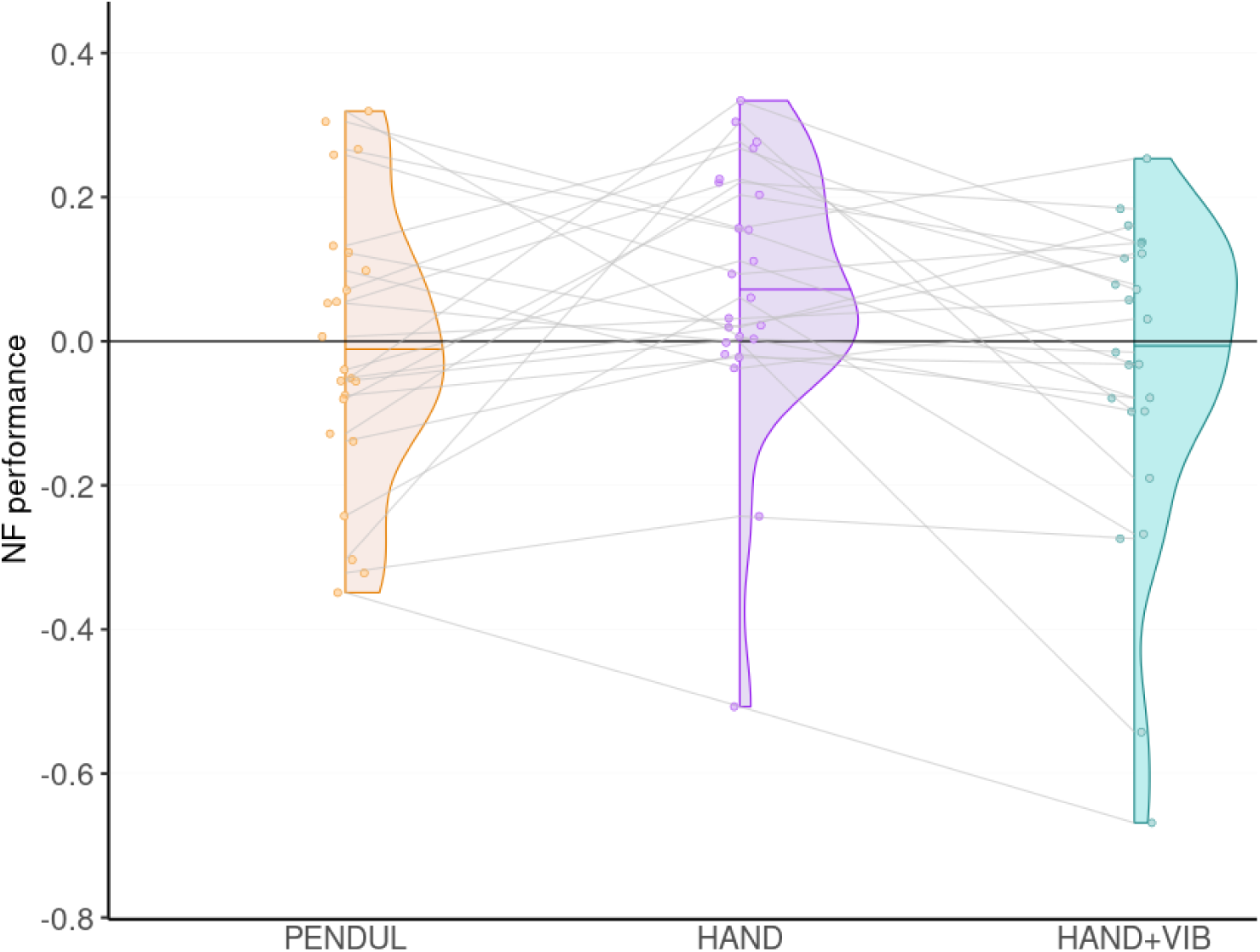
NF performance in the three FB conditions (PENDUL, HAND and HAND+VIB). The distribution of NF performance across subjects is represented as a half violin plot in each FB condition. The horizontal thin color line within each plot represents the median NF performance and the individual data are plotted as coloured points.

We then assessed the effect of time on NF performance for each FB condition, within and across trials using a LMER modelling approach.

In all conditions, the first value of NF performance corresponded to the initial 1.5s of the trial where participants performed MI without any FB yet and clearly stood out as much lower than the other values (Figure 4). It was therefore excluded from the analysis. The NF performance then remained relatively stable across the rest of the trial.

**Figure 4.**
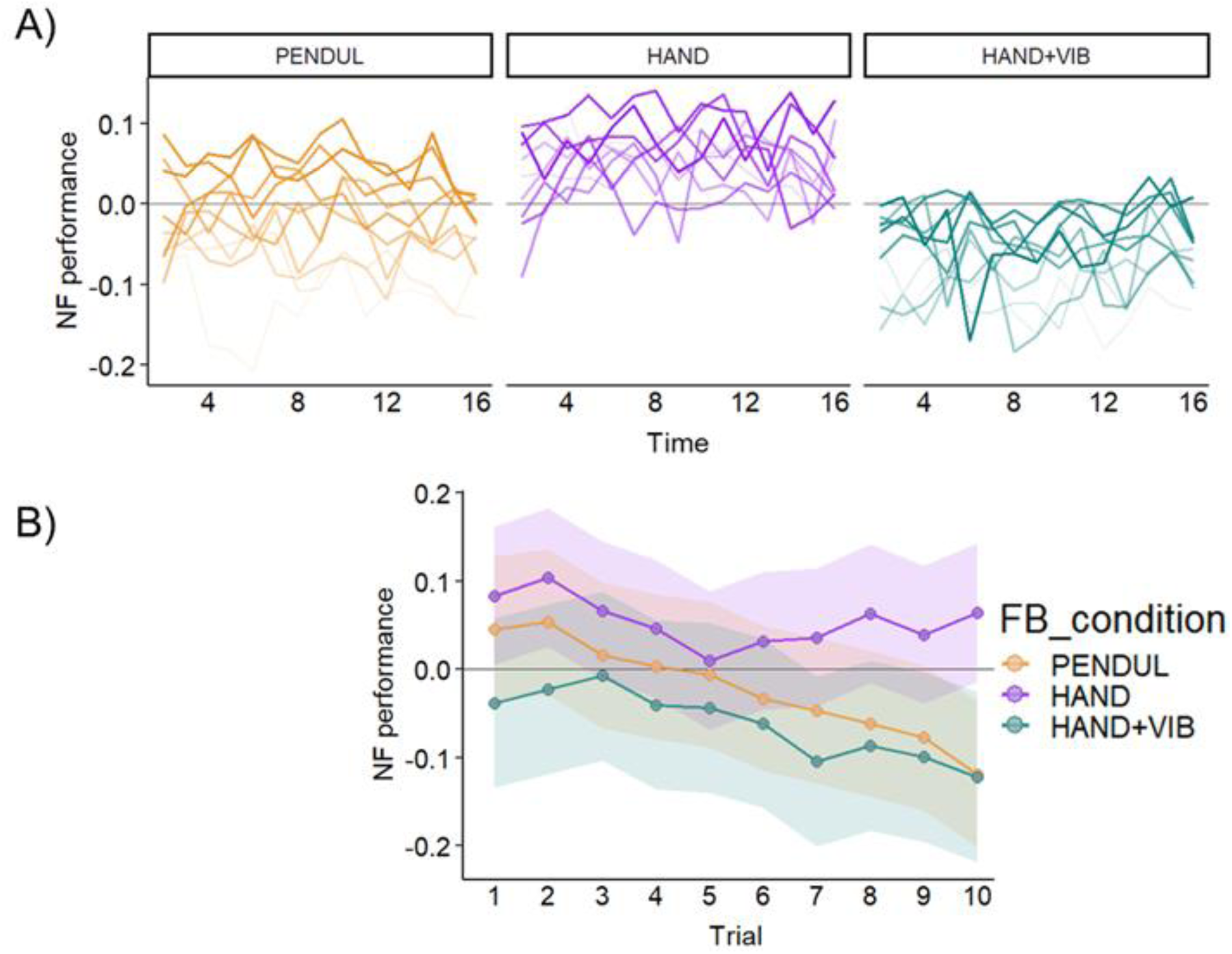
NF performance across the series of FB movements and trials in the three FB conditions. ***A***. Evolution of NF performance across the 16 cycles of FB movement (in abscissa) for each trial, in each FB condition (left to right: PENDUL, HAND, HAND+VIB). The mean of NF performance across the 23 participants is presented. The trial sequence from 1 to 10 is colour coded by colour intensity, with more intense colours for the first trials in the sequence. ***B*.** The same data are presented averaged across cycles to visualise NF performance changes across trials in each FB condition. Points represent the means of the data across subjects. Lines represent the predictions of the model with Trial coded as a categorical factor and shaded areas represent 95% confidence intervals around the model estimates.

Moreover, when examined across trials, NF performance appeared to rapidly degrade for both PENDUL and HAND+VIB. In contrast, HAND FB appeared to uphold NF performance across trials (see Figure 4A, Supplementary Table 2). The LMER model analysis showed main effects of FB_conditions (F(2, 22) = 4.27, p = .027) and Trials (F(9, 10253) = 30.22, p < .0001) and a significant interaction between Trials and FB_conditions on NF performance (F(18, 10253) = 6.69, p < .0001; Figure 4B). This reflected that the differences between HAND and the other two conditions (PENDUL, HAND+VIB) increased along Trials (see Supplementary Table 3 for post-hoc pairwise comparisons of conditions on each trial).

We further tested the linear trend of the evolution of NF performance across trials by computing the LMER model with Trials coded as a numerical factor. This analysis confirmed the interaction between Trials and FB_conditions (F(2, 10277) = 33.67, p < 0.0001), indicating that this interaction was at least partly due to different linear trends of NF performance across the three FB conditions. The linear trend was largest for PENDUL (slope estimate for Trials (est.): -0.018; 95% confidence interval (CI): [-0.020 -0.016]), followed by HAND+VIB (est.: -0.011; CI: [-0.014 -0.009]). In contrast, the trend was an order of magnitude smaller for HAND (est.: -0.004; CI: [-0.006 -0.002], Supplementary Table 4). The pairwise differences in linear trends of NF performance among FB conditions were significant (all p < .0005; see Supplementary Table 5). This confirmed different dynamics of NF performance across trials between FB conditions.

We performed a set of control analyses. First, we checked that the order of presentation of the FB conditions did not significantly impact NF performance (F<1). Second, we determined the potential influence of EOG and EMG artefacts. The results supported that the differences in NF performance between conditions were neither related to EOG nor EMG artefacts (see Supplementary section IV). Third, we tested if the effect of FB conditions on NF performance could be driven by the visual and tactile stimuli themselves. To do so, we tested if these stimuli influenced β power, and thus the calculated performance, during passive conditions, where the participant observed random movements of the pendulum, of the virtual hand, or received vibrations every 4 cycles of the randomly moving virtual hand. Note that for the sake of this analysis, we computed offline performance for the control conditions in the same way as during NF. The ANOVA with FB_conditions and Task (Passive / NF) showed a significant main effect of Task (F(1, 20) = 18.74, p<0.001, η^2^= .26), reflecting lower performance (i.e., stronger β power) during passive than the NF tasks (M = -0.17, SD = 0.29) (Figure 5A and B). The main effect of FB_conditions was not significant (F(2, 40)=1.41, p = 0.25, η^2^= .01). The interaction between Task and FB_conditions was significant (F(2, 40) = 5.70, p = .007, η^2^= .05). This reflected that during the passive task, vibrations were associated with greater offline performance (M = - 0.10, SD = 0.25) than the virtual hand was (M = -0.21, SD = 0.30; t(20) = 2.99, p = 0.007, d = 0.65). Thus, if anything, HAND+VIB should have fostered better NF performance than HAND condition, but we found the opposite effect. There was no difference between passive observation of the virtual hand and pendulum (M = -0.20, SD = 0.31; t(20) = 0.24, p = 0.82, d = 0.05). Finally, we also tested the effect of MI alone. The one-way ANOVA with Task (NF / passive / MI alone) as factor showed a significant main effect of Task (F(2, 38) = 13.48, p = <0.001, η^2^= .42), reflecting higher performance not only in NF (M = 0.02, SD = 0.15) relative to the passive tasks (M = -0.16, SD = 0.27; as shown by the previous analysis) but also in NF relative to MI alone (M = -0.09, SD = 0.23; t(19) = 3.13, p = 0.006, d = 0.70) (Figure 5C). MI alone was associated with greater offline performance (i.e., reduced β power) as compared to the passive tasks (t(19) = 2.88; p = 0.01; d = 0.64).

**Figure 5.**
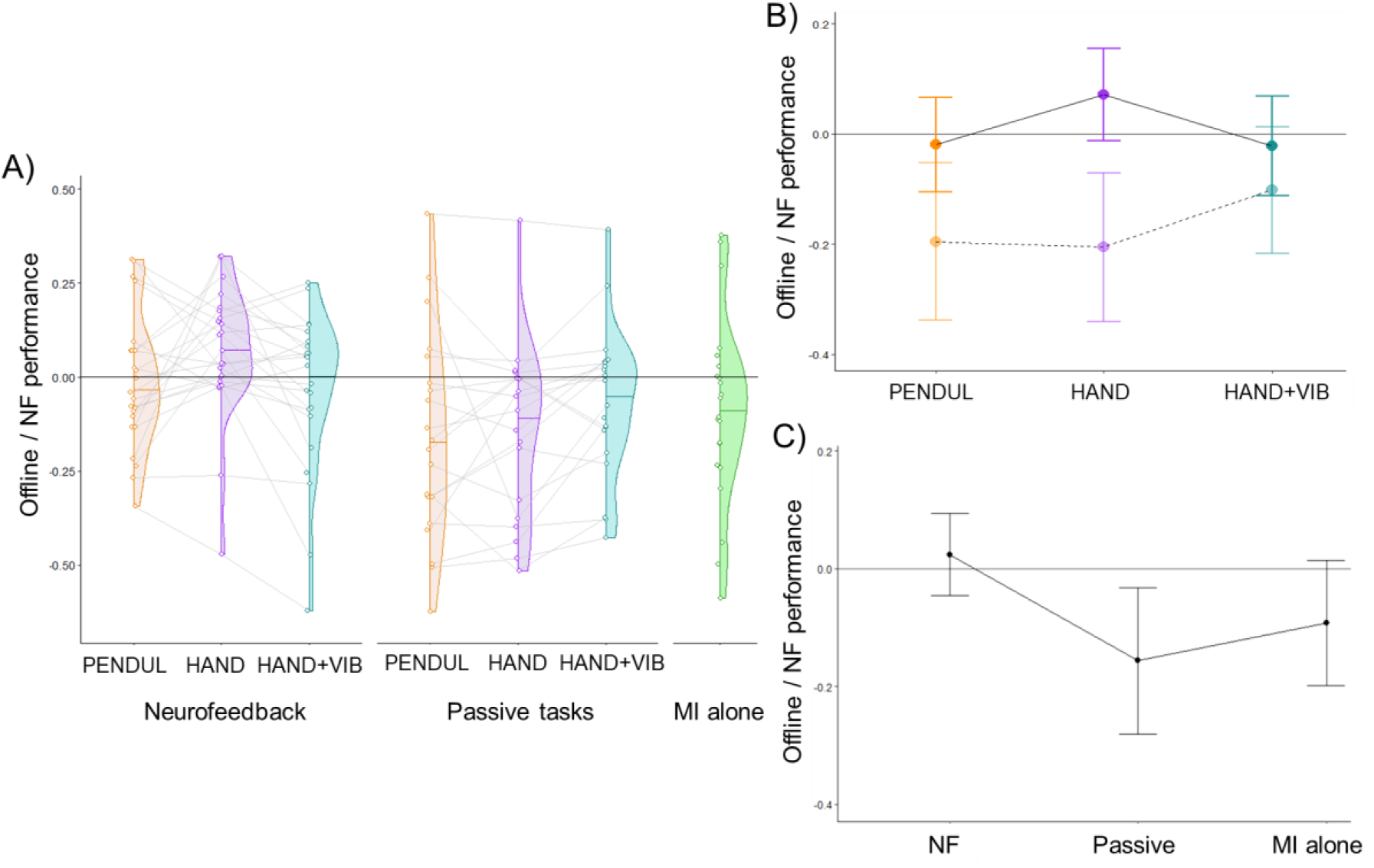
Offline / NF performance comparison in the NF, passive, and MI alone tasks. Offline performance reflected β power modulations during the passive and MI alone tasks. ***A***. NF / offline performance in each condition of NF, passive and MI alone tasks is represented as half violin plots of data distribution with the horizontal line corresponding to the median across subjects and individual data as coloured points. ***B***. Mean NF/ offline performance across subjects for the NF and passive tasks as a function of FB stimulus conditions. The NF tasks are represented in full colours and solid lines; the passive tasks are represented in transparent colours and dotted lines. The vertical bars represent 95% confidence intervals of the means. ***C.*** Comparison of the mean NF / offline performance across subjects in the NF, passive and MI alone tasks. The mean of the 3 stimulus conditions (PENDUL, HAND, and HAND+VIB) is presented for the NF and passive tasks. The vertical bars represent the 95% confidence intervals of the means.

### 3.3 Does transparency increase agency? (H2)

We tested the effect of FB conditions on the sense of agency, which was measured after each run using questionnaires (see Methods; Supplementary Table 6). The 2-way ANOVA with FB_conditions and Run as within-subject factors revealed a significant main effect of FB_conditions (F(2,44) = 4.36, p = .02, η^2^ = .12). This reflected significantly higher agency levels when training with HAND (M = 5.8, SD = 2.5) than PENDUL (M = 4.4, SD = 2.9; t(22) =1.93; p =0.033; d = 0.4). In line with NF performance results, agency was lower in HAND+VIB (M = 4.02, SD = 2.6) relative to HAND (t(22) = 2.7; p = 0.01; d = 0.57) (Figure 6). There was neither any significant main effect of Run (F(1, 22) = 2.23, p > 0.15) nor interaction between FB_conditions and Run (F(2, 44) = 1.69, p > 0.2).

**Figure 6.**
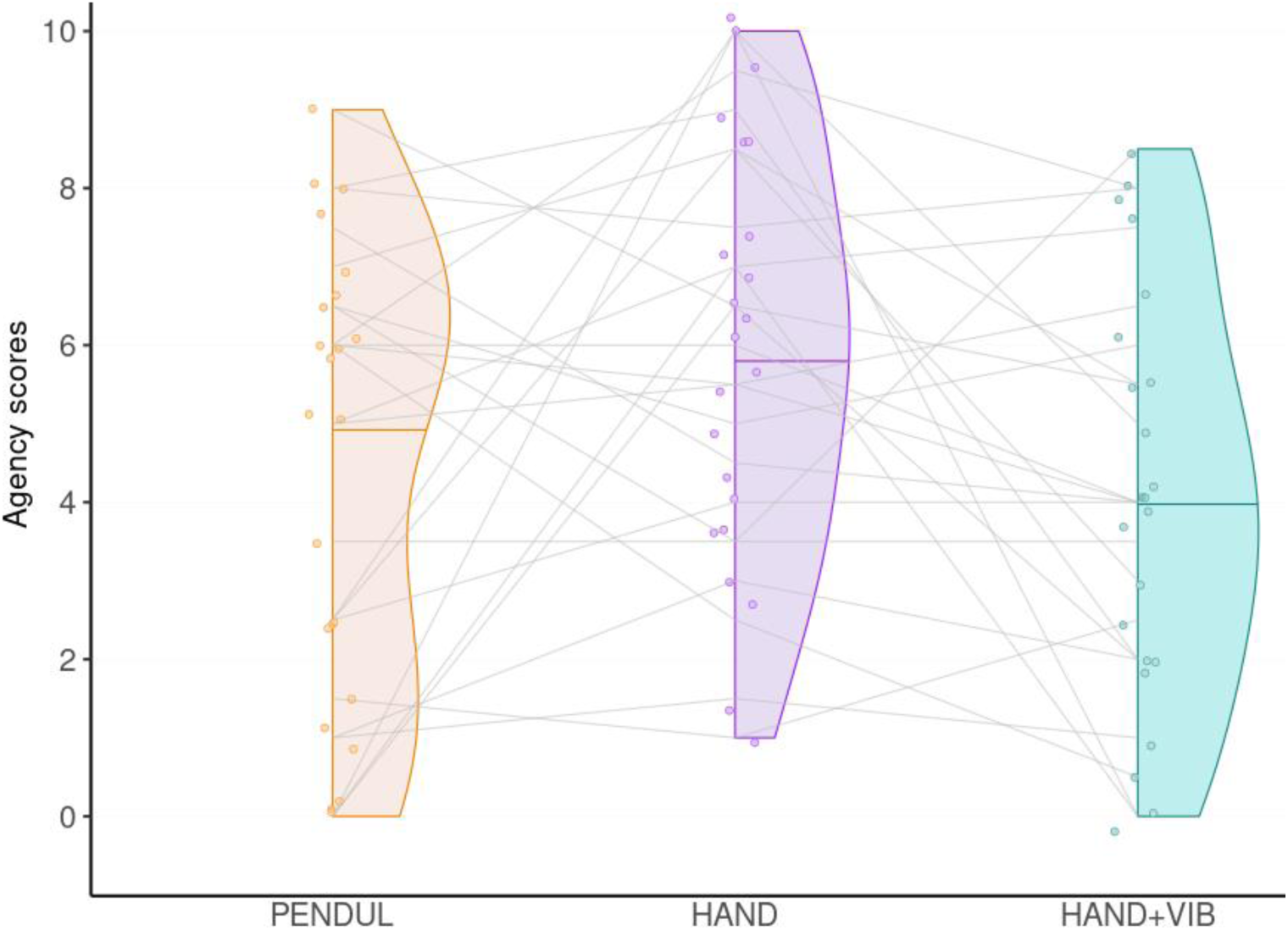
Agency judgements for the three FB conditions (PENDUL, HAND, and HAND+VIB). The distribution of agency scores (averaged across runs) is represented as a half violin plot for each FB condition. The horizontal thin coloured line within each plot represents the median agency score and the coloured points represent the individual data.

Did FB conditions influence other measured variables in relation to the participants’ experience? First, we explored if the increase in agency for the HAND could be related to an overall perception of increased control, whether external or internal. Thus, we tested if the FB conditions were associated with different levels of external causal attribution (Supplementary Table 6). We did not find any effect of FB_conditions (F < 1). External causal attribution scores decreased across runs (F(1, 22) = 4.48, p = 0.05, η^2^ = .04). There was no interaction between FB_conditions and Run (F<1) (Figure 7A). Then, we analysed the effect of FB conditions (restricted to HAND and HAND+VIB) on ownership. The mean ownership judgement was greater for HAND (M = 4.7, SD = 2.8) than HAND+VIB (M = 3.4, SD = 2.8), but this effect was not significant (F(1,44) = 3.3, p = .06, η^2^ = .12). There was neither a main effect of Run (p>0.42) nor an interaction between FB_condition and Run (p > 0.27) on ownership judgement (Figure 7B). For the user*_*experience scores, the ANOVA did not reveal any significant effect (all p > 0.2) (Figure 7C and Supplementary Figure 6). As for feedback_usability, HAND was on average easier to use (M = 0.61, SD = 1.64) than PENDUL (M = -0.31, SD = 1.93) and HAND+VIB (M = -0.30, SD = 1.99), but this effect was not significant (F(2,44) = 2.93, p = .06, η^2^ = .12) (Figure 7D and Supplementary Figure 7). In sum, FB conditions had a significant effect on agency only.

**Figure 7.**
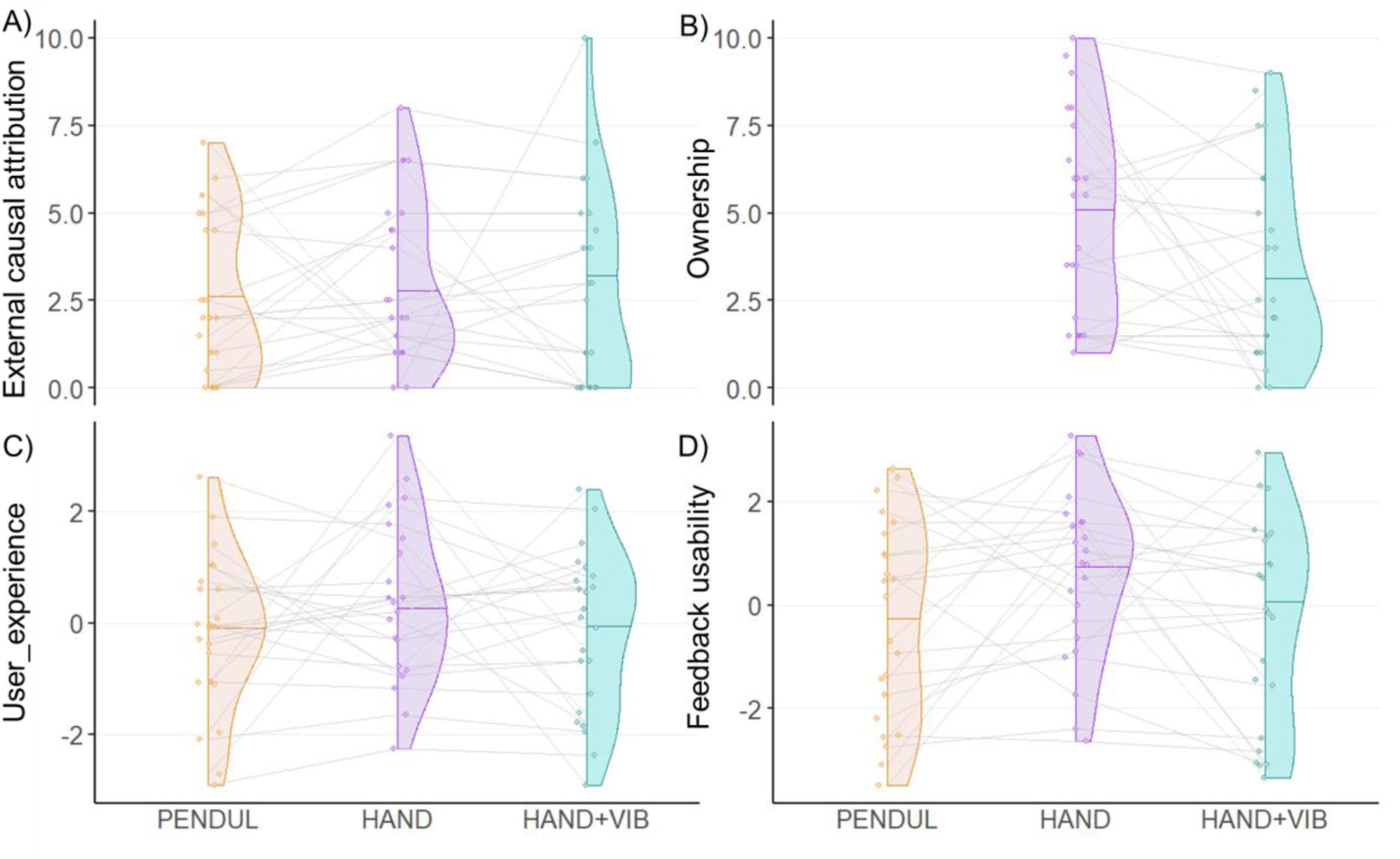
Scores for control questionnaire items in the three FB conditions (PENDUL, HAND, and HAND+VIB). A. External causal attribution. B. Ownership. C. User_experience. D. Feedback_usability. Ownership was measured for HAND and HAND+VIB only. In each plot, the distribution of the individual scores (averaged across runs in A, B and C) is represented as a half violin plot for each FB condition. The horizontal thin coloured line within each plot represents the median score and the coloured points represent the individual data.

### 3.4 Agency captures a large part of the variance related to the effect of FB transparency on NF performance (H3)

FB condition appeared to modulate both NF performance and agency in the same direction (i.e., higher NF performance and agency in HAND than in PENDUL and HAND+VIB). We went on testing if these effects were related, with the hypothesis that agency may capture a large part of the variance of the effect of FB transparency on NF performance. To test this hypothesis, we used incremental LMER models with NF performance as outcome variable and FB conditions and/or agency as predictor variables. The model with only FB_conditions and Run as fixed effects showed a significant main effect of FB_conditions (Model 1: F(2, 24.66) = 5.38, p = 0.012, see Table 2). Including Agency as an additional fixed effect (Model 2) yielded a significant main effect of Agency (F(1, 21.87) = 16.7, p = 0.0005) while the effect of FB_conditions was rendered insignificant (Model 2: F < 1, see Table 2). Model comparison showed that Model 2 outperformed Model 1 (χ²(1) = 13.2, p = 0.0003; see Table 2, Model 1 vs. Model 2 comparison). Moreover, Model 2 did not outperform a more parsimonious model including only Agency and Run as fixed effects (Model 3) (χ²(2) = 1.33, p = 0.51; see Table 2, Model 2 vs. Model 3 comparison).

**Table 2.**
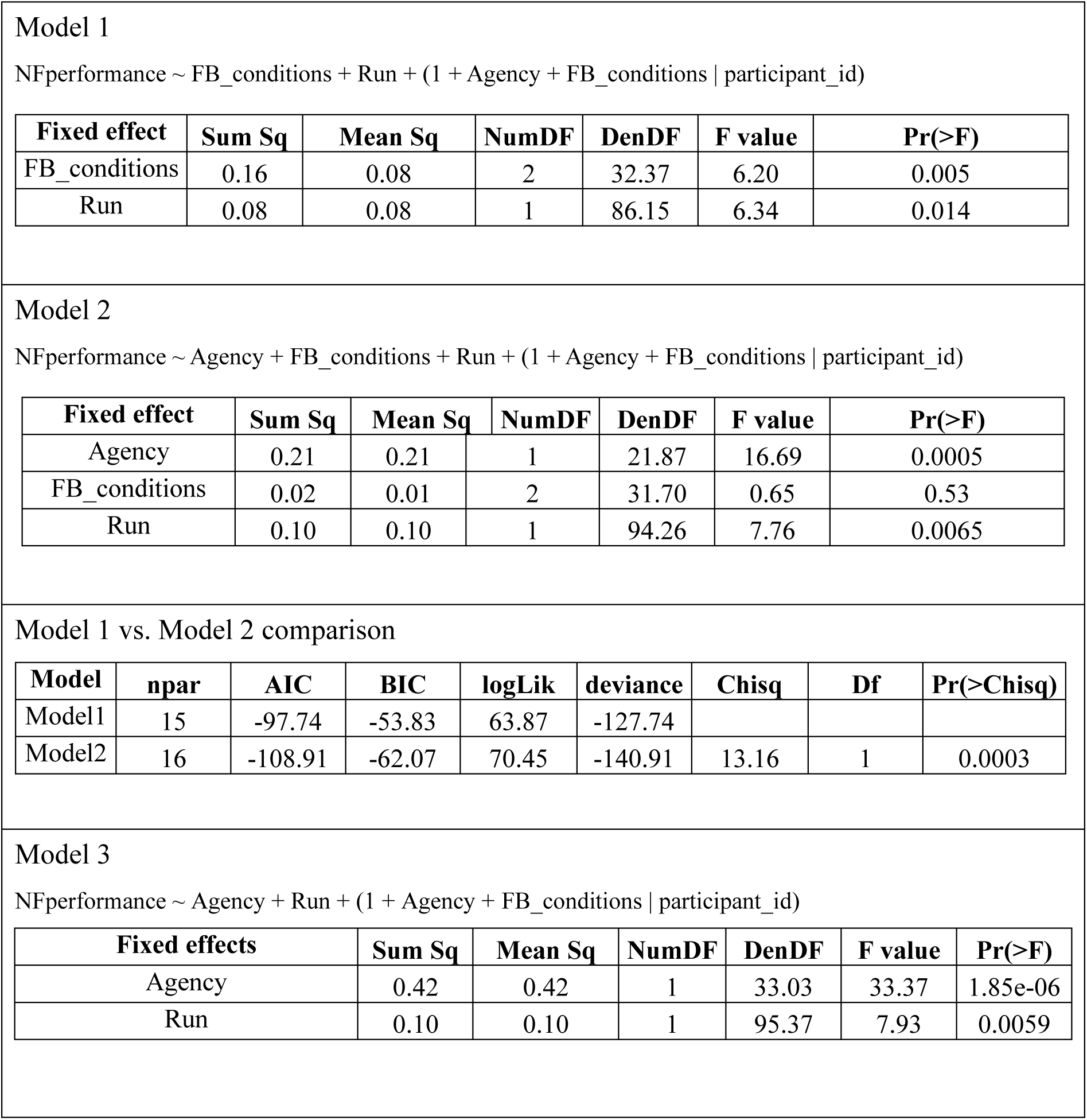
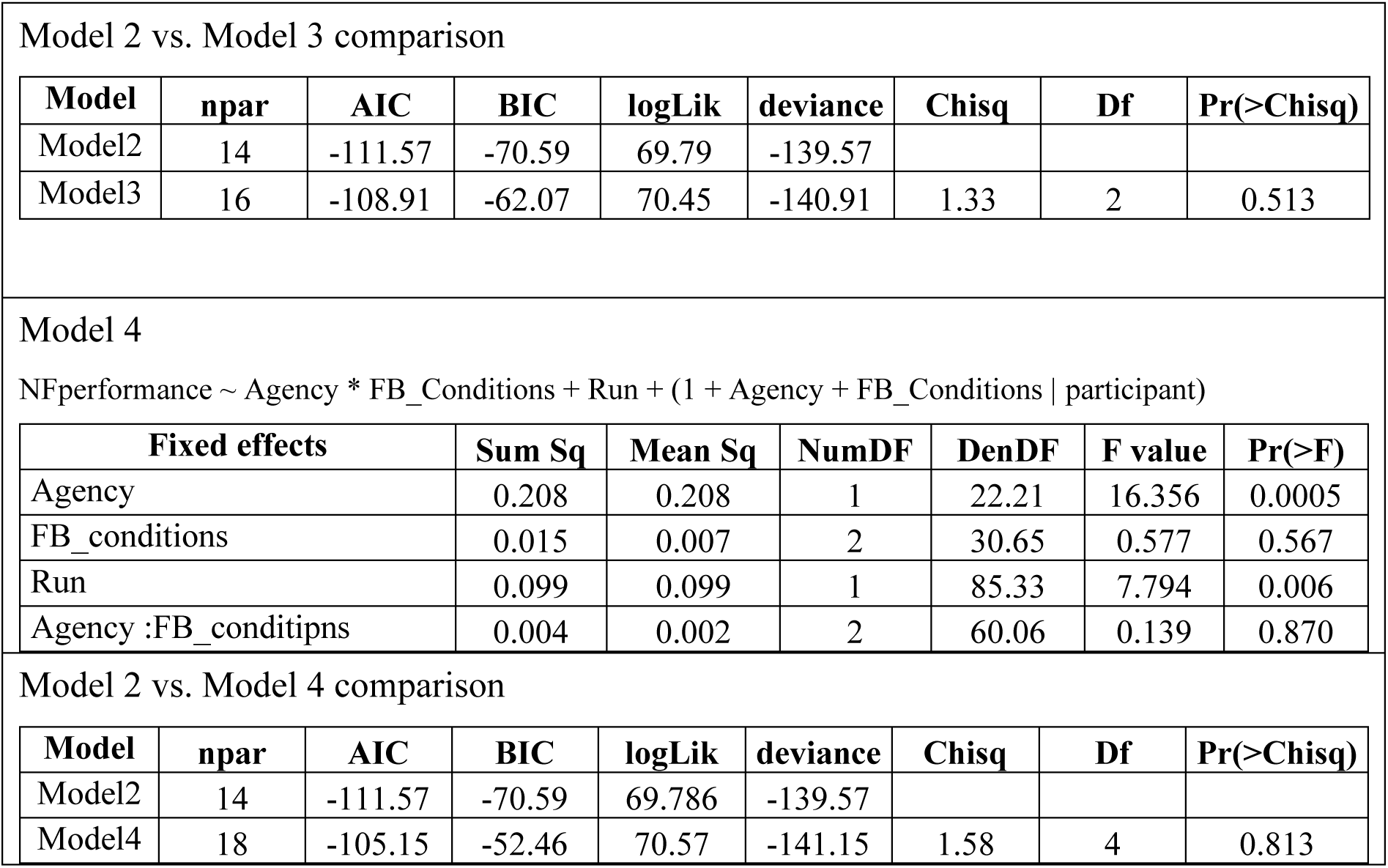
Results of the incremental LMER models of NF performance, testing the contributions of FB_conditions and Agency.

| Fixed effect | Sum Sq | Mean Sq | NumDF | DenDF | F value | Pr(>F) |
| --- | --- | --- | --- | --- | --- | --- |
| FB_conditions | 0.16 | 0.08 | 2 | 32.37 | 6.20 | 0.005 |
| Run | 0.08 | 0.08 | 1 | 86.15 | 6.34 | 0.014 |

Model 2 NFperformance ~ Agency + FB\_conditions + Run + (1 + Agency + FB\_conditions | participant\_id)
| Fixed effect | Sum Sq | Mean Sq | NumDF | DenDF | F value | Pr(>F) |
| --- | --- | --- | --- | --- | --- | --- |
| Agency | 0.21 | 0.21 | 1 | 21.87 | 16.69 | 0.0005 |
| FB_conditions | 0.02 | 0.01 | 2 | 31.70 | 0.65 | 0.53 |
| Run | 0.10 | 0.10 | 1 | 94.26 | 7.76 | 0.0065 |

Model 1 vs. Model 2 comparison
| Model | npar | AIC | BIC | logLik | deviance | Chisq | Df | Pr(>Chisq) |
| --- | --- | --- | --- | --- | --- | --- | --- | --- |
| Model1 | 15 | -97.74 | -53.83 | 63.87 | -127.74 |  |  |  |
| Model2 | 16 | -108.91 | -62.07 | 70.45 | -140.91 | 13.16 | 1 | 0.0003 |

Model 3 NFperformance ~ Agency + Run + (1 + Agency + FB\_conditions | participant\_id)
| Fixed effects |  | Sum Sq | Mean Sq | NumDF | DenDF | F value | Pr(>F) |
| --- | --- | --- | --- | --- | --- | --- | --- |
| Agency |  | 0.42 | 0.42 | 1 | 33.03 | 33.37 | 1.85e-06 |
| Run |  | 0.10 | 0.10 | 1 | 95.37 | 7.93 | 0.0059 |

Model 2 vs. Model 3 comparison
| Model | npar | AIC | BIC | logLik | deviance | Chisq | Df | Pr(>Chisq) |
| --- | --- | --- | --- | --- | --- | --- | --- | --- |
| Model2 | 14 | -111.57 | -70.59 | 69.79 | -139.57 |  |  |  |
| Model3 | 16 | -108.91 | -62.07 | 70.45 | -140.91 | 1.33 | 2 | 0.513 |

Model 4 NFperformance ~ Agency \* FB\_Conditions + Run + (1 + Agency + FB\_Conditions | participant)
| Fixed effects | Sum Sq | Mean Sq | NumDF | DenDF | F value | Pr(>F) |
| --- | --- | --- | --- | --- | --- | --- |
| Agency | 0.208 | 0.208 | 1 | 22.21 | 16.356 | 0.0005 |
| FB_conditions | 0.015 | 0.007 | 2 | 30.65 | 0.577 | 0.567 |
| Run | 0.099 | 0.099 | 1 | 85.33 | 7.794 | 0.006 |
| Agency :FB_conditipns | 0.004 | 0.002 | 2 | 60.06 | 0.139 | 0.870 |

Model 2 vs. Model 4 comparison
| Model | npar | AIC | BIC | logLik | deviance | Chisq | Df | Pr(>Chisq) |
| --- | --- | --- | --- | --- | --- | --- | --- | --- |
| Model2 | 14 | -111.57 | -70.59 | 69.786 | -139.57 |  |  |  |
| Model4 | 18 | -105.15 | -52.46 | 70.57 | -141.15 | 1.58 | 4 | 0.813 |

Finally, introducing the interaction between Agency and FB_conditions in the LMER model (Model 4) did not improve model fit (p = 0.81, see Table 2, Model 2 vs. Model 4 comparison) and Model 4 showed no significant interaction between Agency and FB_conditions (Model 4: F < 1; p = 0.87). We illustrate the relation between Agency and NF performance in Figure 8.

**Figure 8.**
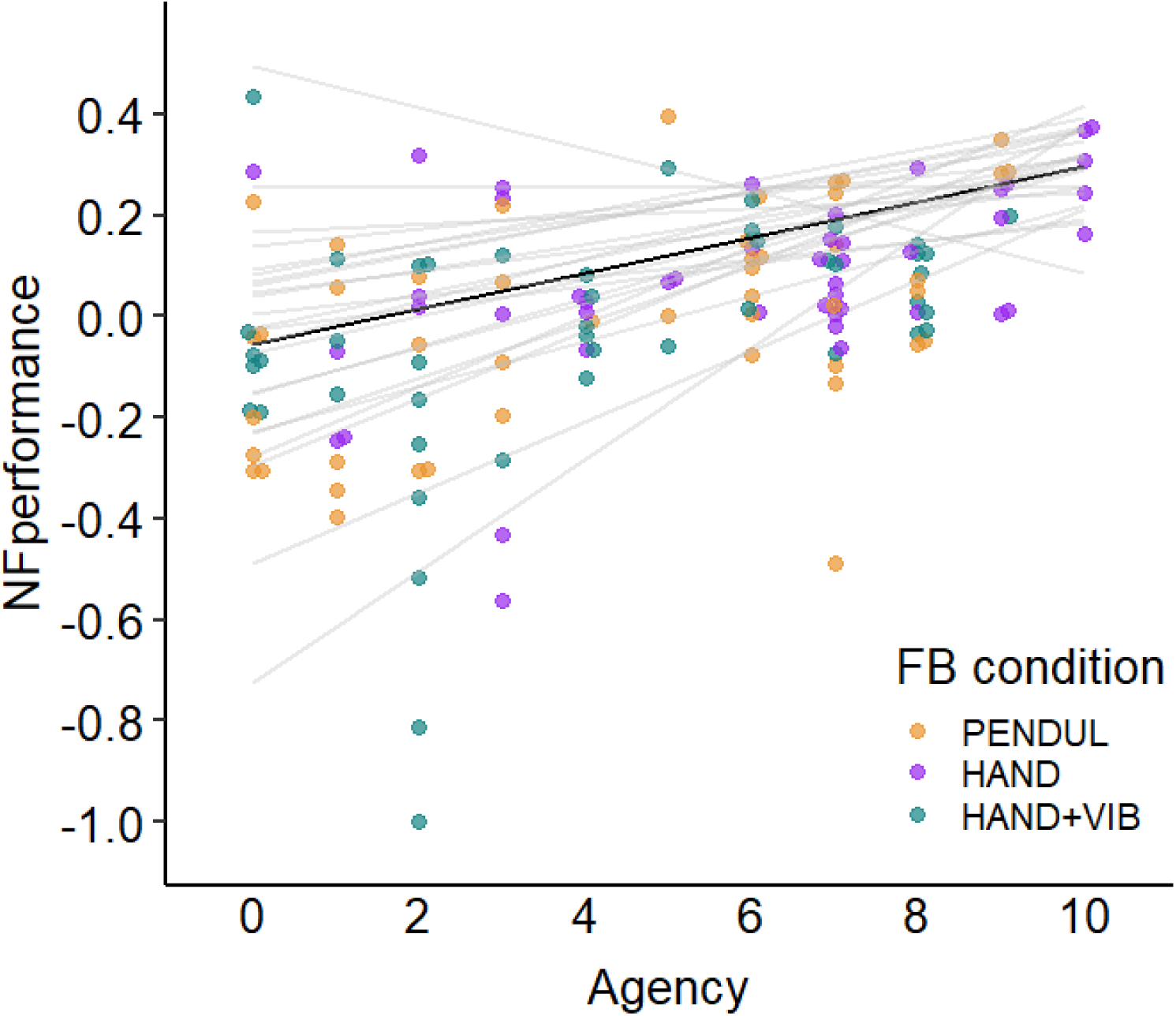
Relation between agency and NF performance. NF performance (y-axis) is plotted as a function of agency (x-axis). Individual data are represented as filled dots of a different colour for each FB condition (PENDUL in orange, HAND in purple and HAND+VIB in turquoise), with one data point for each run for every subject. The thin gray lines represent the individual random slopes and intercepts of the effect of agency on NF performance, as estimated from Model 2. The black thick line represents the estimated fixed effect of agency.

In sum, the incremental LMER analyses showed that agency fully captured the variance related to the effect of FB conditions on NF performance. There was a positive relation between agency and NF performance (parameter estimate for the Agency effect in Model 2 = 0.04, 95% CI [0.02, 0.05]; F(1, 21.87) = 16.69, p = 0.0005; Table 2) and this relation did not depend on the FB condition.

## 4. Discussion

This study investigated the effect of FB transparency on NF performance, with the hypothesis that FB transparency would increase NF performance and that this effect would be related to agency. We targeted β band regulation in a MI-EEG NF protocol and used a pendulum (PENDUL), a virtual hand (HAND), and a virtual hand with vibrations inducing motor illusion (HAND+VIB) as FB of assumed increasing transparency. Time-frequency analysis over the scalp showed the expected 8-30 Hz desynchronisation pattern, peaking on left central electrode C3 in all FB conditions. We found an effect of transparency on NF performance for HAND relative to PENDUL, but contrary to our hypothesis, the addition of intermittent vibrotactile stimuli in the HAND+VIB condition had a detrimental effect on performance. Passive control conditions indicated that this was not related to the sensory properties of the FB conditions. Furthermore, the effect of FB conditions on NF performance varied across trials, with a steady linear degradation of NF performance for the PENDUL and HAND+VIB conditions, which was significantly reduced for the HAND condition.

Importantly, FB conditions were associated with different agency ratings, with higher agency for the HAND than for both PENDUL and HAND+VIB, mirroring the effect of FB on NF performance. We further analysed the link between NF performance, FB conditions and agency. We showed a positive relation between agency and NF performance and agency modulation fully captured the variance of the effect of FB conditions on NF performance.

### 4.1 H1 - Transparency improves NF performance

We assumed increased transparency across our 3 FB conditions (PENDUL < HAND < HAND+VIB). This postulate was only partly verified. The rhythmically opening and closing virtual hand seemed indeed more transparent than the swaying pendulum because it reproduced the movement that the participant was imagining. In line with our hypothesis, this resulted in increased NF performance, corresponding to greater *online β* power reduction, for HAND relative to PENDUL.

To our knowledge, only one previous NF study compared an abstract bar FB to a virtual hand FB (Ono et al., 2013). These authors reported no difference in online NF performance between their FB conditions. This may be because they manipulated FB transparency between-subjects, with only eight participants for each FB type. In contrast, we manipulated FB transparency in a within-subject manner, with twenty-three participants analysed in our three FB conditions. This allowed us to demonstrate the effect of visual transparency on NF performance.

Other studies investigated BCI performance differences when training with abstract bar versus realistic virtual hand FB in between-subjects designs. These paradigms trained participants to perform left and right MI, thus performance related to the accuracy in differentiating between those two tasks. Neuper et al. (2009) reported no difference in online classifier accuracy when comparing an abstract visual bar to a virtual hand FB in a between-subject study. They concluded that the presentation form of FB does not impact BCI performance when the FB informational content is equivalent. In the present study, the informational content of PENDUL and HAND was matched with identical mapping of *online β* power to PENDUL and HAND quantity of movement. Yet, we found an effect of transparency with better NF performance for HAND than PENDUL. While this does not exclude a contribution of informational content in some previous studies, it supports the idea that the transparency of visual FB plays a role in NF performance. In the same line, Skola & Liarokapis. (2018) and Penaloza et al. (2018) reported better BCI performance with virtual hands than bar FB. Moreover, our study included several control conditions, including a passive condition of FB stimulus observation. This allowed us to rule out a confounding effect of action observation since there was no difference in *online β* power (aka. ‘NF performance’) between PENDUL and HAND in this condition.

Contrary to our initial hypothesis, the addition of vibrotactile FB inducing motor illusion did not improve but rather decreased performance in comparison to the HAND condition. This contradicts studies suggesting that adding motor illusions to a virtual hand FB can improve MI BCI performance (Yao et al., 2015; Barsotti et al., 2018), which inspired us to include this condition (see also Jeunet et al., 2015). We think that this discrepancy arose from some of our FB design choices, which may have reduced the transparency of the visuo-tactile FB relative to the visual FB. First, the vibrations were delivered intermittently (every 6 s), to avoid habituation and movement illusions in the opposite direction (Taylor et al., 2017). As a result, the vibratory FB was infrequent, rendering it potentially distracting. Second, the vibrotactile FB was an all-or-nothing, binary FB, in contrast to the continuous visual FB. Third, the tactile FB was computed based on a longer time window than the visual FB. As a result, visual and vibrotactile FB were decoupled and participants received vibrations without seeing hand movements in about a third of cases (Supplementary Figure 3). This mismatch between the visual and tactile modalities in the HAND+VIB condition may have rendered it less transparent than the HAND FB. This interpretation is supported by the fact that the HAND FB was overall judged as easier to use than the other conditions.

It is interesting to note that during the passive control condition, the addition of vibrations in HAND+VIB condition reduced *online β* power as compared to HAND condition. Thus, if anything, the vibrotactile FB should have been beneficial to NF performance. In agreement with this idea, one can note that ERD seemed stronger in the time intervals of vibratory FB in Supplementary Figure 5. This is in line with the studies that showed contralateral ERD in the mu & beta band in response to motor illusion vibrations (Yao et al., 2015; Schneider et al., 2021). Furthermore, providing motor illusion vibrations during MI allowed for stronger ERD than MI alone did (Le Franc et al., 2021). However, these authors noted that the ERD in response to passive vibrations was not different from the ERD during the MI with vibrations condition. Thus, the effect they observed was likely driven by the vibrations. In sum, based on their intrinsic effect, one may have expected the vibrations to facilitate NF performance. This reinforces our idea that the design of our HAND+VIB FB condition may have actually reduced transparency, resulting in decreased NF performance relative to the HAND condition. Importantly, taken together with the lack of difference in *online β* power (aka. ‘NF performance’) for HAND and PENDUL in the passive control condition, this supports the idea that the sensory properties of our FB stimuli did not account for the differences in NF performance.

Further analyses of NF performance evolution across trials revealed distinct trends for the HAND versus the other FB conditions. The NF performance was maintained along trials for HAND in comparison to both PENDUL and HAND+VIB. Performance degradation across trials in the PENDUL and HAND+VIB conditions may reflect a progressive disengagement of the participants from the task, related to the lower overall subjective usability of the less transparent FB. This degradation was observed over a single session of NF, emphasizing the importance of FB transparency. Refining FB design may improve usability and agency, fostering increased task engagement (Jeunet et al., 2016), which may result in improved NF performance, thus supporting responsiveness to NF.

### 4.2 H2 – FB transparency improves sense of agency

The HAND condition was associated with higher agency ratings than the PENDUL and the HAND+VIB conditions. Agency is based on three key principles: priority (temporal contiguity of action intention, action, and outcome), consistency (matching of predicted and sensory outcome), and exclusivity (uniqueness of the apparent cause of the outcome) (Wegner & Wheatley, 1999). Action-FB correspondence is key to establishing agency, irrespective of the form of the effector (Caspar et al., 2015; Zopf et al., 2018). Violation of consistency and priority reduces sense of agency over VR virtual hands (Jeunet et al., 2018). Action-FB correspondence relates to FB transparency in our design. To the best of our knowledge, the first study that linked these concepts to agency used a fake BCI, providing positive reward independent of the user’s brain activity (Beursken, 2012; Van Acken, 2012). By comparing virtual hands that either matched the imagined movement (transparent FB) or did not (thumb-up FB provided during fist clenching MI), these authors found that transparent FB was associated with a higher sense of control over the FB, akin to agency.

The impact of task-FB transparency on agency was explored by BCI studies with real FB in two ways. A set of BCI studies manipulated FB congruency, i.e., FB direction relative to the imagined movement, for example displaying left movement in response to right (resp. left) hand MI in incongruent (resp. congruent) condition. Using either abstract (cursor) (Evans et al., 2015; Marchesotti et al., 2017) or transparent virtual or robotic hand FB (Braun et al., 2016; Serino et al., 2022), these studies reported higher sense of agency with congruent than incongruent FB. Notably, Evans et al’s (2015) study featured a random FB condition; they showed that this false FB elicited agency when it was congruent with the imagined movement. Moreover, a study manipulated task-FB transparency by varying the subject’s task: the participants were required to control a virtual arm FB with either an MI (transparent condition) or a SSVEP-based NF protocol. This study reported greater agency for the MI task that matched with the FB (Nierula et al., 2021). Our result brings additional evidence that transparency is key to agency.

Transparency did not influence significantly the other variables measured in relation to the participants’ experience (external causal attribution, ownership, user_experience, feedback_usability). External causal attribution was a low-level control, because the participants always tried to action the FB in our study. Yet, it yielded a significant effect of run but no significant effect of FB conditions. Regarding ownership, it was measured for HAND and HAND+VIB conditions only and did not show any significant difference between the two FB conditions. It is possible that the sense of ownership contributed to improved agency for HAND versus PENDUL feedback, since we did not assess ownership for PENDUL. In this regard, prior work manipulating ownership levels might provide some insight. Alimardani et al. (2016b) manipulated ownership by comparing humanoid robotic hands to rudimentary metallic grippers as FB. Similarly, Ziadeh et al. (2021) compared virtual hands to abstract blocks. In both studies, the two conditions had equivalent task-FB movement mapping and participants showed equivalent agency levels between the two conditions but different levels of ownership. This suggests that ownership is not sufficient to foster agency and thus may not explain our results. Finally, the effects of FB conditions on user_experience and on feedback_usability were not significant, although the HAND FB was judged on average easier to use than the other FB.

Adding a motor illusion to the HAND FB reduced participants’ sense of agency. Pillette et al. (2021a) showed higher agency over visuo-tactile FB than visual-only virtual hand FB, which suggests that it is unlikely that the tactile FB modality itself caused the agency degradation. However, visuo-tactile synchrony is key to inducing agency over virtual hands as shown in studies using the rubber-hand illusion (Tsakiris et al., 2006; Ratcliffe & Newport, 2017). In our study, the vibratory FB was computed based using a long time window, which may have reduced the temporal contiguity between action intention and outcome, a key component for the sense of agency. Every participant but one experienced incongruency between the vibrotactile and the visual FB in the form of tactile FB without virtual hand FB movement (Supplementary Figure 3). Thus, our finding of reduced agency for the HAND+VIB condition is consistent with the results of Alimardani et al. (2016a) who showed that a multimodal FB condition featuring temporally mismatched visual and proprioceptive signals leads to a reduced sense of agency compared to a visual-only FB condition (see also Hanashima & Ohyama, 2022). Moreover, by manipulating the congruency of visual and tactile FB with unimodal and multimodal FB conditions, Serino et al. (2022) have recently shown that agency is more affected by tactile than visual FB incongruency. This may suggest that visual-only FB is preferable to visuo-tactile FB with mismatched sensory inputs (Alimardani et al., 2016a; Caspar et al., 2021).

### 4.3 H3 - Agency captures a large part of the variance related to the effect of transparency on NF performance

NF performance and agency showed a similar pattern of variation with FB conditions. When considered together, agency fully captured the variance related to the effect of FB conditions on NF performance. This highlighted a strong, likely bidirectional, link between agency and NF performance, independent of FB.

Our study is the first to investigate the link between performance and agency in a NF paradigm. Such link was investigated in BCI paradigms with mixed results. Most of these studies conceptualized performance as a basis for agency rather than agency as a lever for performance. This aligns with research outside the BCI domain, which showed agency correlated with perceived performance even when performance was influenced by external factors outside of the participant’s control (Metcalfe & Greene, 2007). In the BCI domain, a study featuring virtual hand FB (Ziadeh et al., 2021) found a correlation between BCI performance and agency (see also Pillette et al, 2021b). Two other studies reported no correlation between BCI performance (Skola & Liarokapis, 2022) or motor alpha ERD (Nierula et al., 2021) and sense of agency in MI-based paradigms. Some studies manipulated FB-task congruency, presenting participants with false FB independent on their brain activity and making this FB either congruent (perceived as positive performance) or incongruent (perceived as negative performance) relative to the MI cue. With this paradigm, Evans et al (2015) showed that congruent trials led to higher agency than incongruent ones did, suggesting that a false sense of agency can arise from perceived BCI performance regardless of actual brain activity modulation. It is possible that their FB manipulation disrupted the consistency principle of agency in an extreme way, since their incongruent FB was in opposite direction to the imagined movement, thus featuring inversed mapping of FB and MI task. In contrast, with a similar paradigm, Marchesotti et al (2017) found a positive correlation between agency and performance in true-FB trials but a negative one in manipulated congruent/incongruent trials.

In our study, we found a robust link between NF performance and agency. This link is likely to be bidirectional. e. On one hand, as discussed above, better NF performance may have reinforced agency and feeling of control. On the other hand, agency, as influenced here by FB transparency, may have fostered NF performance. This suggests that leveraging FB design to increase agency could be key to initiating a virtuous circle that fosters motivation, engagement, and reinforcement learning, which are key to NF learning (Strehl, 2014; Gaume et al, 2016). In this line, recent NF studies have begun exploring the manipulation of sense of agency by targeting its correlates in central theta activity (Zito et al., 2023), arguing towards the validity of an actionable loop between sense of agency and neural rhythm modulation.

### 4.4 Limitations

Our work has several limitations. First, as discussed above, our HAND+VIB condition induced a detrimental effect. However, this finding may not be conclusive regarding the effect of transparency due to the visuo-tactile FB mismatch, followed by an expected decline in agency.

We did not explicitly manipulate agency in our design but rather evaluated it after each NF run. Thus, we cannot make inferences about the role of agency as a causal basis for NF performance and the link uncovered here is likely bidirectional. It will be interesting to manipulate sense of agency experimentally in future studies to investigate this issue. Such manipulations have been successfully described outside of BCI paradigms. For instance, virtual hand can be modified to break agency principles (Jeunet et al., 2018). Buccholz et al (2019) designed a visuo-motor task with a no-agency, a hidden agency and an overt agency. Adapting these manipulations to MI-NF protocols could allow investigating the direction of the agency-NF performance relationship.

We evaluated agency after each run of 5 trials. Measuring it after each trial could have allowed finer-grained results. However, it could also have undermined NF performance by requiring frequent task interruptions. Though not tested in a BCI environment, interruptions are known to be detrimental to attention and performance (Bailey & Konstan, 2006), especially in cognitive tasks (Lee & Duffy, 2014). It may also be noted that most previous studies measured the sense of agency only once, at the end of the training (Alimardani et al., 2016a; Braun et al., 2016; Skola & Liarokapis, 2018; Nierula et al., 2021; Ziadeh et al., 2021; Pillette et al., 2021b; Caspar et al., 2021). The few studies that sampled agency after each trial used binary measures (Evans et al., 2015; Marchesotti et al., 2017; Serino et al., 2022), which gives less range of expression to the participant and are generally reported to be less reliable (Markon et al., 2011). In contrast, our design allowed us to sample agency over six measures and to run linear mixed effect model analysis to examine the link between agency and NF performance. Most interestingly, a recent study used the error-related negativity component with EEG to disentangle internal and external causal attribution of virtual hand movements (Gomez-Andres et al, 2024). This could provide within-trial, continuous markers of sense of agency allowing for robust tracking of the association between NF performance and agency.

## 5. Conclusion

Our results show that transparency influences NF performance and agency and they highlight a strong link between agency and NF performance across all FB types. More precisely, we found that transparent visual FB provided by a virtual hand yields better NF performance than abstract FB such as a pendulum. On the other hand, the design of multimodal FB should be carefully considered to provide fully congruent, integrated FB information. Otherwise, detrimental effects may be observed, as was the case here. Importantly, agency was associated with the effect of our FB conditions on NF performance. This highlights a way for improving NF protocols by considering agency and leveraging FB design, which may improve the number of people who are able to learn NF.

## Author contributions

C Dussard, CJK, NG and BL developed the study concept. Study implementation was performed by C Dussard with technical support from LH. The virtual hand was implemented by adapting a previous Unity C# code provided by LP. Data collection was performed by C Dussard with help from N.G for the first participants. C Dussard analysed the data with supervision by N.G, CJK and BL. C Dumas performed artefact analysis with supervision by C Dussard and NG. C Dussard and LP drafted the manuscript. NG, CJK, EP and BL provided critical revisions. All authors critically discussed the results, contributed to their interpretation and provided feedback on the manuscript. All authors approved the final version of the manuscript for submission.

## Acknowledgments

This work was supported by the French National Research Agency (BETAPARK project, ANR-20-CE37-0012). The EEG data were acquired on the CENIR MEG-EEG facility of the ICM, which received funding by the program “Investissements d’avenir” (Agence Nationale de la Recherche, grant numbers ANR-10-IAIHU-06 and ANR-11-INBS-006) for infrastructures.

We thank Aurore Bussalb for her help in the experiments. We thank Véronique Marchand-Pauvert for lending us material. We thank Déborah Ziri for her benevolent proofreading of the final manuscript.

