## Supplementary file for "Influence of feedback transparency on motor imagery neurofeedback performance: the contribution of agency" (from another version of this preprint: version 3, 2024-08-12)

### SUPPLEMENTARY MATERIAL

### I. Reference β level

For some participants, the initially computed *reference β* level (median β power value from 2 30 s baseline runs) was either very low or too high, resulting in either no visual FB at all or constant maximal amplitude movements. Thus, we adjusted the *reference β* level manually at the beginning of the experiment to allow participants to have FB during the task. The initial median values and final values used for each participant are reported in **Supplementary Table 1**. For the first 5 participants (3 included in our analyses), the *reference β* level was computed before each FB condition. Then, for the remaining participants, we decided to keep the *reference β* level constant throughout the experiment to avoid potential confounding effect of baseline variability across conditions.

**Supplementary Table 1. *Reference β* level used for each participant**. The initial value was the median value of the 8-30 Hz C3-Laplacian power during baseline runs. It was adjusted manually to a final value to allow the participant to receive some FB during the task. This value was used throughout the experiment except for the first 5 participants for whom a *reference β* value was computed before each FB condition.

| Participant nb. | Initial value | Final value |
|---|---|---|
| 0 | [20.5, 24.9, 23.6] | [23.8, 20.3, 27.0] |
| 1 (excluded) | [21.3, 25, 40.4] | [15.0, 11.7, 25.0] |
| 2 | [2.9, 2.9, 2.4] | [2.4, 2.8, 2.8] |
| 3 | [20.1, 20.1, 20.1] | [22.3, 22.3, 22.3] |
| 4 (excluded) | [7, 9.5] | [11, 9.6] |
| 5 | 6.4 | 20.0 |
| 6 | 13.5 | 35.0 |
| 7 | 25.9 | 35.0 |
| 8 | 22.3 | 32.0 |
| 9 | 25.3 | 30.0 |
| 10 | 20.6 | 35.0 |
| 11 | 310.8 | 32.0 |
| 12 | 5.8 | 8.5 |
| 13 | 18 | 20.5 |
| 14 | 10.4 | 10.0 |

| | | |
|---|---|---|
| 15 | 11.2 | 20.2 |
| 16 | 10.1 | 15.0 |
| 17 | 7.2 | 10.0 |
| 18 | 13.6 | 13.0 |
| 19 | 9.2 | 13.0 |
| 20 | 8.7 | 17.0 |
| 21 | 16 | 16.0 |
| 22 | 21.1 | 27.5 |
| 23 | 11.4 | 16.0 |
| 24 | 11.3 | 6.0 |

### II. Visual and tactile FB computation and supplementary analysis

#### II.1 Visual and tactile FB computation

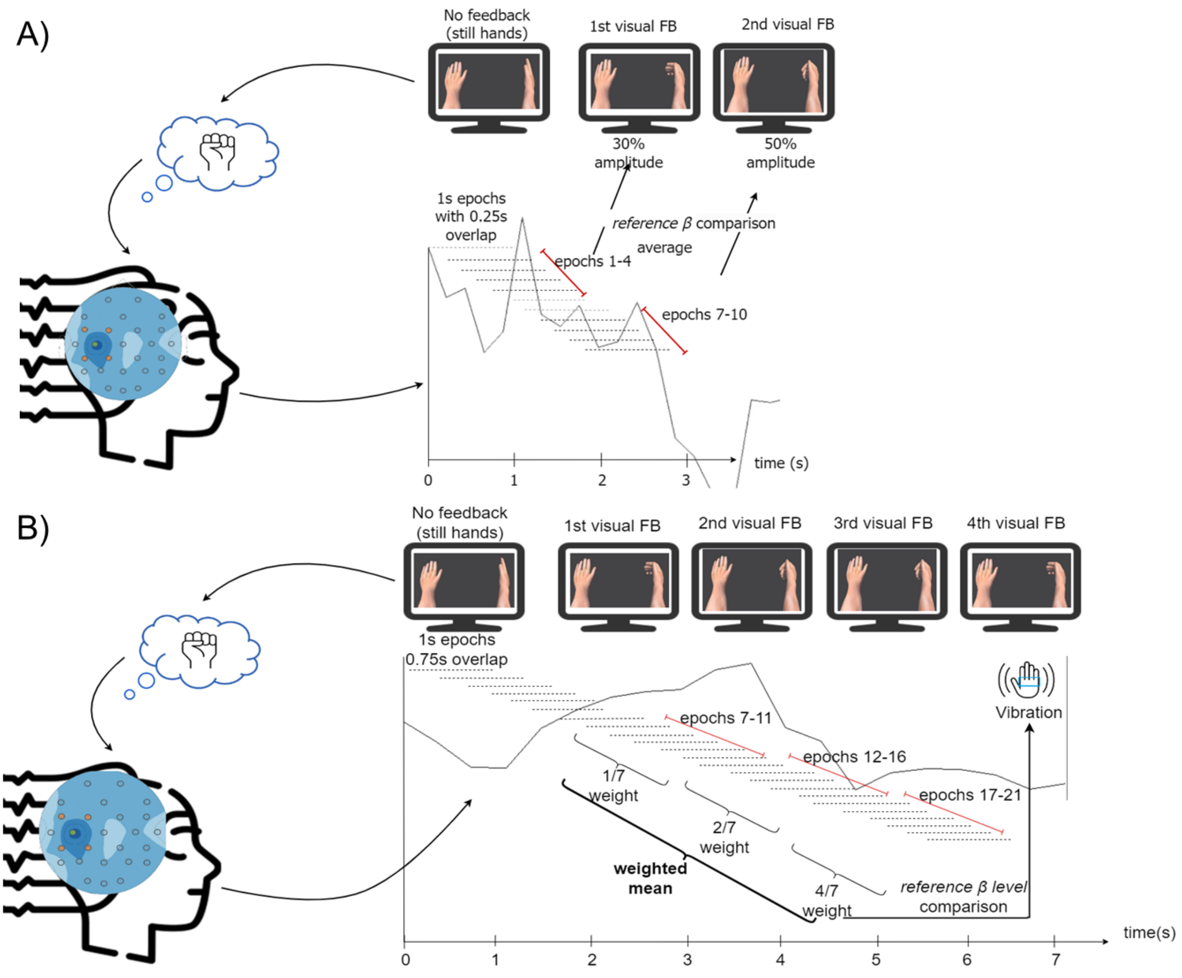


**Supplementary Figure 1.** The *online β* power is represented on the graph as a function of trial time. It was computed in sliding, overlapping windows of 1s, represented as horizontal lines. A. For each cycle of the visual FB movement (here, illustrated for HAND), the *online β* power values of the 4 epochs preceding the movement cycle were averaged and compared to the *reference β* level in order to determine the amplitude of FB movement on the screen. B: For the HAND+VIB FB, which could be delivered on every 4 cycles of HAND FB movement, the mean of the 15 β band power values before the vibration onset was computed in a weighted manner as follows: the 5 values just before the vibration onset were given a weight of 4, the 5 values before were given a weight of 2, and the 5 values furthest away from the vibration onset were given a weight of 1.

### II.2 Supplementary analysis of the amount of visual and tactile FB received

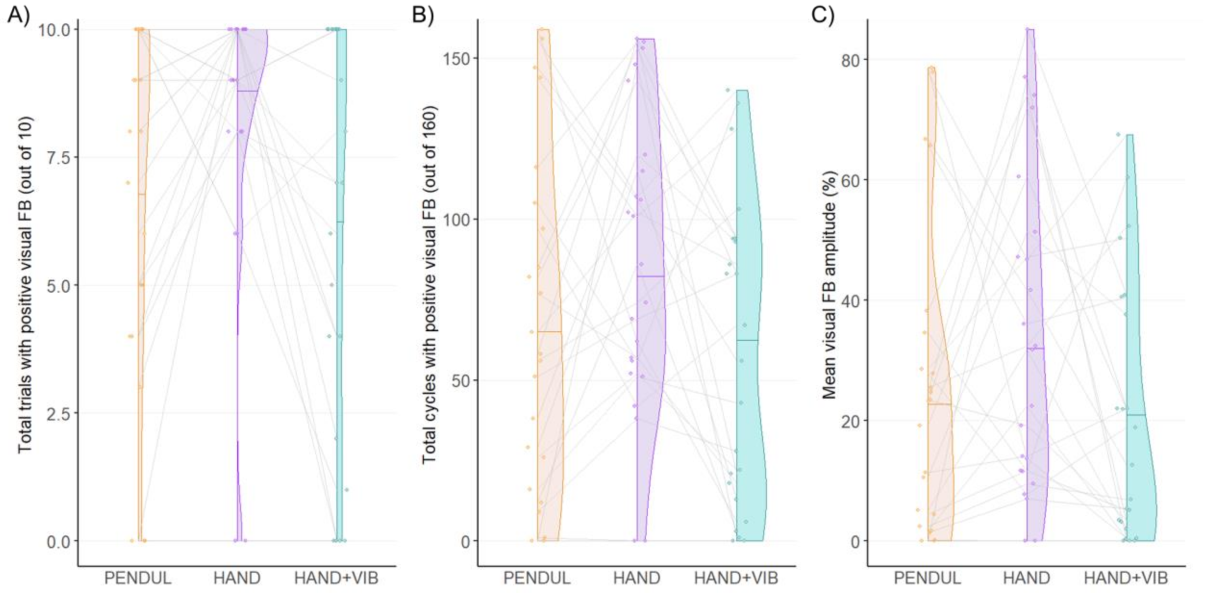


**Supplementary Figure 2.** Amount of FB received by the participants in each FB condition. A. Number of trials with positive visual FB. B. Number of cycles with positive visual FB. C. Mean visual FB amplitude. For each plot, the distribution of values across subjects is represented as a half violin plot for each FB condition. Horizontal thin color lines within each plot represent the median value and the individual data are plotted as coloured points.

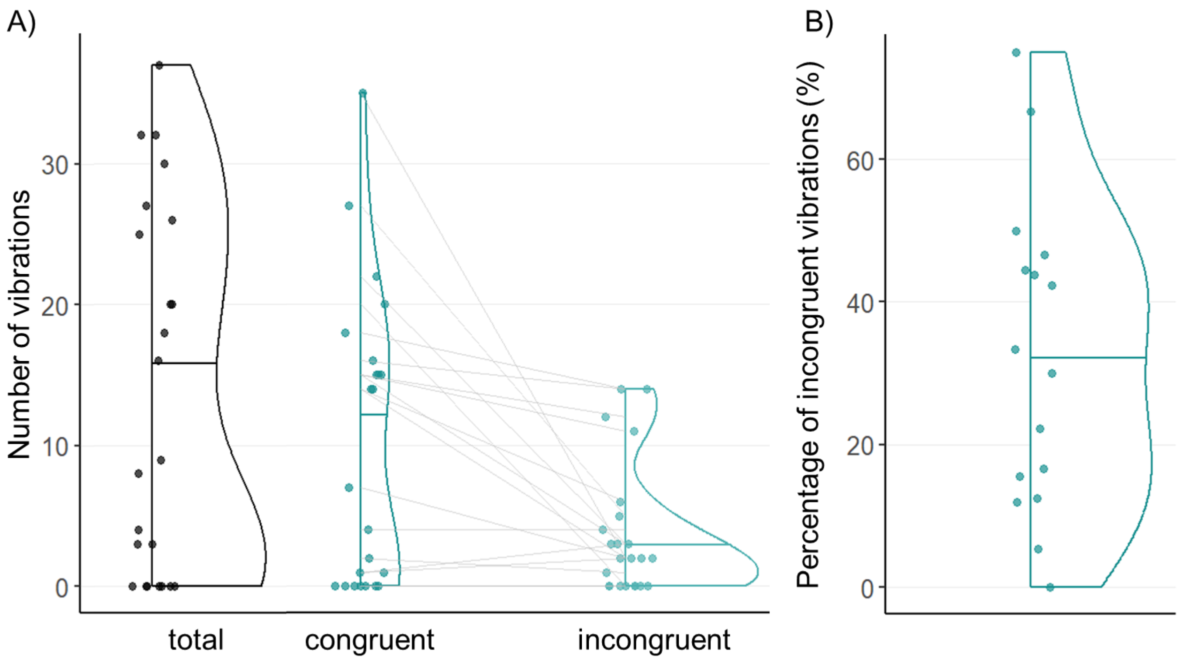


**Supplementary Figure 3.** Number of vibrations received in the HAND+VIB condition. A. Total number of vibrations received (left half violin), number of concomitant tactile and visual FB (“congruent”; middle half violin), and number of tactile FB without any co-occurring virtual hand movement (“incongruent”; right half violin). Total number of vibrations received (left half violin), number of concomitant tactile and visual FB (“congruent”; middle half violin), and number of tactile FB without any co-occurring virtual hand movement (“incongruent”; right half violin). B. Percentage of incongruent vibrations, that is, number of tactile FB without any co-occurring virtual hand movement divided by the total number of vibrations received, times 100. For every plot, the distribution of values across subjects is represented as a half violin plot. Horizontal thin color lines within each plot represent the median value and the individual data are plotted as coloured points.

### III. Time-frequency analysis pipeline

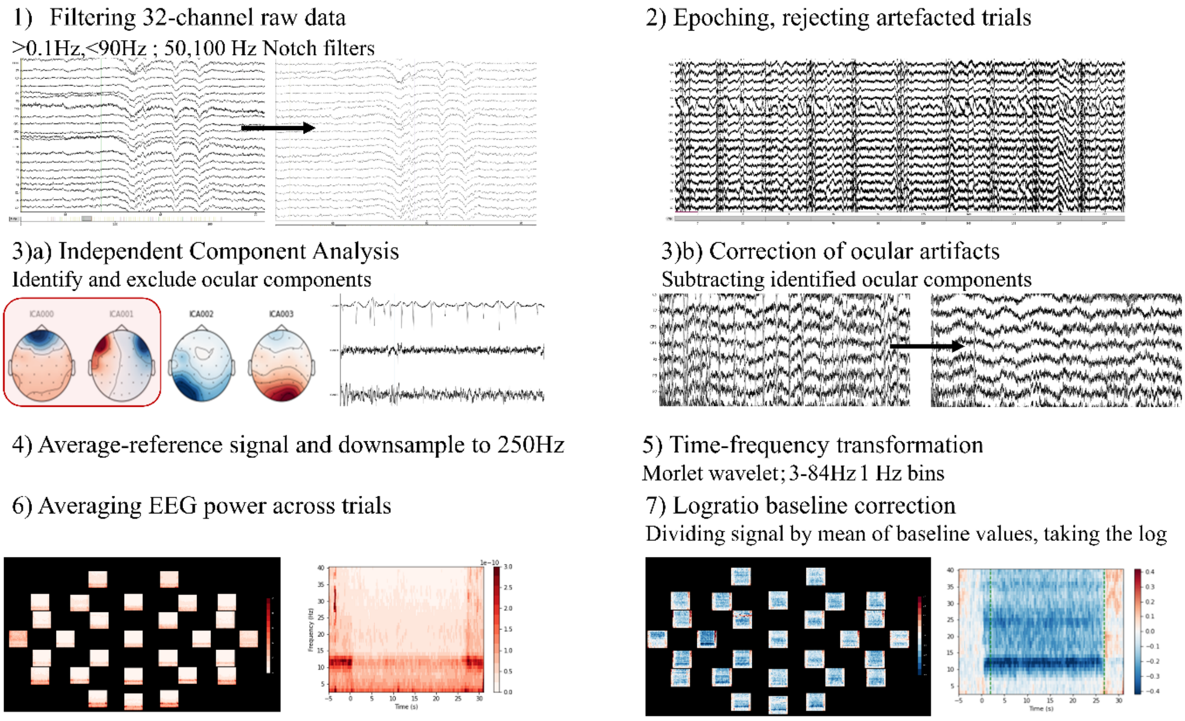


**Supplementary Figure 4. Processing pipeline from raw EEG data to time-frequency representation for one participant and one FB condition**. The successive steps of data processing as detailed in the main text are illustrated. 1) Filtering: the timecourse of example rawdata on each electrode before (on the left) and after (on the right) filtering is represented. 2) Epoching. An exemple time course of epoched data is illustrated. 3a) Independent component analysis (ICA): On the left, illustration of the topographical maps of 4 example ICs, with ICA000 and ICA001 corresponding to blinks and horizontal saccades respectively. On the right, time course of ICA000, ICA001, and ICA002. 3b) Correction of ocular artefact: example time course of data before and after ICA ocular artefact correction on electrodes C3, T7, CP5, CP1, Pz, P3 and P7 . 6) Averaging EEG power across trials: example time-frequency (TF) representation of the data on all electrodes (on the left) and illustration of an example TF map on C3 Electrode. 7) Logratio baseline correction. Same representation as in 6.

### IV. Control analysis of the impact of artefacts on NF performance

We performed artefact detection on NF trials raw data in each FB condition to check if there was a potential effect of artefacts in our results.

*Artefact detection*

Automatic artefact detection was performed using the Python library MNE (version 0.23.0). The signal was filtered between 1 and 10Hz. Blinks were detected using the find_eog_events function with VEOG and HEOG channels. For 6 participants out of the 23, Fp1 and Fp2 were used instead because of poor EOG signal quality.

Muscle artefacts were detected using the annotate_muscle_zscore function with default parameters on T8, T7, FC1, FC5, O1, O2, Oz, CP1, CP5, C3, F7 and F8 channels (adapted from Goncharova et al., 2003).

For both types of artefacts, the MNE annotations were reviewed and corrected manually. Muscle artefacts yielded more variable annotations than blinks, requiring extensive correction of the muscle segments.

For each FB cycle, we considered the four 1s epochs used to compute *online β* power values for this cycle. The cycle was considered artefacted if one of these epochs contained an artefact. This was coded as a factor "blink_presence" or 'muscle_artefact_presence".

*Statistical analyses*

We tested whether the median NF performance was significantly different between artefacted and non artefacted cycles by using a 2-way ANOVA with FB_conditions and artefact_presence as within-subject factors. Only the 9 participants who had muscle artefacts were included in the 2-way ANOVA for EMG artefacts.

*Results*

The 2-way ANOVA revealed no significant effect of blink_presence on median NF performance ($F(1,44) = 0.001$, $p = .97$, $\eta^2 = .000001$) and only replicated the effect of FB_conditions on NF performance ($F(2,44) = 6.09$; $p = 0.007$; $\eta^2 = 0.20$). There was no significant interaction between blink presence and FB_conditions ($F(2,44) = 0.89$, $p = 0.41$, $\eta^2 = 0.002$).

The 2-way ANOVA in the subgroup of participants with muscle artefacts did not show any main effect of either muscle artefact presence ($F(1, 16) = 1.97$; $p = 0.20$, $\eta^2 = 0.009$ ) or FB_conditions ($F(2, 16) = 0.88$, $p = 0.39$; $\eta^2 = 0.09$) on median NF performance. There was no significant interaction between muscle_artefact_presence and FB_conditions ($F (2, 16) = 0.19$, $p = 0.80$, $\eta^2 = 0.001$).

At the scale of our study, our analysis reflected that the difference in NF performance across FB conditions did not seem driven by blinks or muscle artefacts.

### V. Time-frequency representation of EEG power at C3 in each FB condition.

PENDUL HAND HAND+VIB

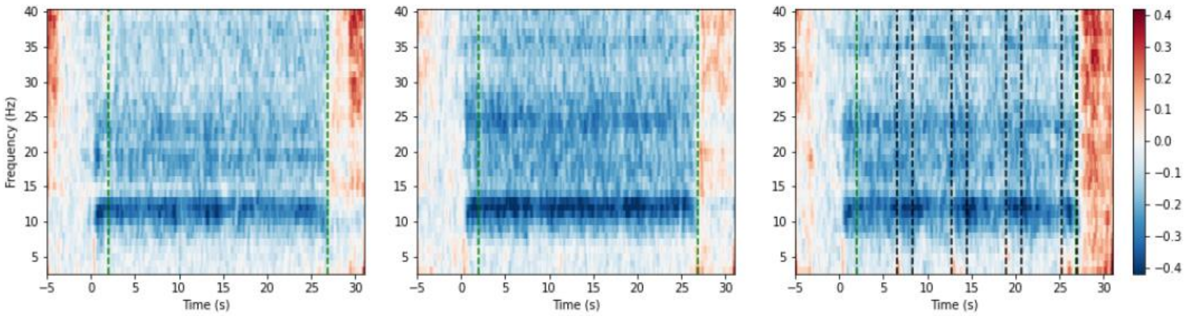


**Supplementary Figure 5. Time-frequency representation of EEG power at C3 in each FB condition.** Time-frequency data are represented between 3 and 40 Hz (in ordinate) along the trial time (in s, in abscissa). The data were averaged across the NF trials for each FB conditions. The grand average across the 23 participants is shown for—from left to right— PENDUL, HAND, and HAND+VIB. The green dashed lines represent the start and end of FB movement cycles. The black dashed lines represent vibration periods in the HAND+VIB condition. ERD/ERS values are color coded, with the blue colors representing ERD and the red colors representing ERS (in dB).

### VI. Additional details on statistical analyses.

The LMER model structure was determined through nested testing. An interaction between FB_conditions and Trials was observed upon visual data examination (Figure 4A), which was confirmed by model comparison (anova function of stats R package): Including the 2-way interaction between FB_conditions and Trials accounted significantly better for the data than the additive model alone, as reflected by Akaike Information Criterion (AIC) and Bayesian Information Criterion (BIC). In contrast, Time appeared to have only an additive effect and model comparison showed a non-significant contribution of the two-way interaction between FB condition and Time. Therefore, the latter interaction term was not included in the final model.

**Supplementary Table 2. ANOVA table of the LMER model used for NF performance analysis, with Trials coded as a categorical factor**

Type III Analysis of Variance Table with Satterthwaite's method

| | Sum Sq | Mean Sq | NumDF | DenDF | F value | Pr(>F) |
|---|---|---|---|---|---|---|
| FB_conditions | 0.36 | 0.18 | 2 | 22 | 4.27 | 0.027 * |
| Trials_factor | 11.36 | 1.26 | 9 | 10253 | 30.22 | < 2e-16 *** |
| Time | 0.01 | 0.01 | 1 | 10253 | 0.20 | 0.65 |
| FB_conditions:Trials_factor | 5.03 | 0.28 | 18 | 10253 | 6.69 | < 2e-16 *** |

Significance codes: '***' 0.001; '**' 0.01; '*' 0.05; '.' 0.1

The post-hoc comparisons were performed with the emmeans package, using the following command lines:

emmeans(model,~FB_conditions |Trials,at=list(FB_conditions=c(PENDUL,HAND)))

emmeans(model,~FB_conditions |Trials,at=list(FB_conditions=c(HAND,HAND+VIB)))

Where model refers to the LMER model used for NF performance analysis as specified in the main text of Method, with Trials coded as a categorical factor.

**Supplementary Table 3. Post-hoc comparison of NF performance for PENDUL vs. HAND and HAND vs. HAND+VIB conditions, on every trial.**

| Trial\ FB comparison | PENDUL vs. HAND | HAND+VIB vs.HAND |
|---|---|---|
| **1** | z = 0.82, p = 0.46 | **z = 2.8, p =0.008** |
| **2** | z = 1.09, p = 0.39 | **z = 2.94, p = 0.007** |
| **3** | z = 1.1, p = 0.39 | z = 1.72, p = 0.1 |
| **4** | z = 0.93, p = 0.44 | z = 2.00, p = 0.06 |
| **5** | z = 0.34, p = 0.73 | z = 1.25, p = 0.21 |
| **6** | z = 1.41, p = 0.32 | **z = 2.16, p = 0.04** |
| **7** | z = 1.80, p = 0.18 | **z = 3.26, p = 0.003** |
| **8** | **z = 2.71, p = 0.03** | **z = 3.48, p = 0.003** |
| **9** | **z = 2.54, p = 0.04** | **z = 3.21, p = 0.003** |
| **10** | **z = 3.99 p < 0.01** | **z = 4.32, p = 0.002** |

The z and p values of the post-hoc tests are indicated. P values are corrected with FDR (false discovery rate) correction for each column (10 tests corresponding to each emmeans command)

As can be seen in the Supplementary Table 3 the difference between HAND and PENDUL conditions was initially statistically non-significant and then increased, with a significant difference in trials 8, 9 and 10. For HAND vs HAND+VIB conditions, the difference was statistically significant in all trials but trials 3, 4 and 5.

We also performed post-hoc analyses with the emtrends function fo the emmeans package, using the following command line: emtrends(model,"FB_conditions",var=Trials"). Here, model referred to the LMER model used for NF performance analysis as specified in the main text of Method, with Trials coded as a numerical factor.

**Supplementary Table 4. Post-hoc analysis of the slope trends of NF performance along trials for each FB condition**

| FB_conditions | Trials.trend | SE | df | asymp.LCL | asymp.UCL |
|---|---|---|---|---|---|
| HAND | 0.004 | 0.0012 | Inf | 0.002 | 0.006 |
| HAND+VIB | 0.012 | 0.0012 | Inf | 0.009 | 0.014 |
| PENDUL | 0.018 | 0.0012 | Inf | 0.016 | 0.020 |

Trials.trend: estimate of the linear trend across trials for each FB condition (in rows).

Degrees-of-freedom method: asymptotic.

Asymp.LCL/UCL = asymptotic lower and upper (resp.) confidence levels, with confidence level = 0.95

**Supplementary Table 5. Pairwise comparisons of the linear trends of NF performance along trials, between FB conditions**

| contrast | estimate | SE | df | z.ratio | p.value |
|---|---|---|---|---|---|
| HAND - PENDUL | -0.014 | 0.002 | Inf | -8.20 | <.0001 |
| HAND – HAND+VIB | -0.008 | 0.002 | Inf | -4.41 | <.0001 |
| HAND+VIB - PENDUL | -0.007 | 0.002 | Inf | -3.79 | 0.0004 |

Analysis performed with the pairs function applied to the emtrends computed in the post-hoc analysis of Supplementary table 5.

Degrees-of-freedom method: asymptotic

P value adjustment: Tukey method for comparing a family of 3 estimates

### VII. Questionnaires

**Supplementary Table 6. Median scores and interquartile ranges (IQR) of questionnaires in each FB condition across runs.**

| Questionnaire item | **PENDUL** | | **HAND** | | **HAND+VIB** | | Phase |
|---|---|---|---|---|---|---|---|
| | Median | IQR | Median | IQR | Median | IQR | |
| *Agency self* | 5.5 | 5.75 | 7 | 4.75 | 4 | 5 | Post-run |
| *External causal attribution* | 2 | 4 | 2 | 4.75 | 2 | 4 | Post-run |
| *Embodiment* | _ | _ | 4 | 6 | 2.5 | 4 | Post-run |
| *Success* | 5 | 5 | 5 | 4.75 | 5 | 5 | Post-run |
| *Difficulty* | 6.5 | 2.75 | 6 | 4 | 6.5 | 3 | Post-run |
| *Satisfaction* | 5 | 3 | 5 | 2.75 | 5 | 3 | Post-run |
| *Easiness* | 6 | 5.5 | 7 | 2 | 6 | 4.5 | Post-FB |
| *Reliability* | 6 | 4 | 6 | 4 | 6 | 3.5 | Post-FB |
| *Learnability* | 6 | 6 | 7 | 3.5 | 5 | 5 | Post-FB |
| *Esthetics* | 7 | 2.5 | 8 | 3 | 7 | 3 | Post-FB |
| *Reactivity* | 5 | 4.5 | 6 | 3 | 5 | 5.5 | Post-FB |

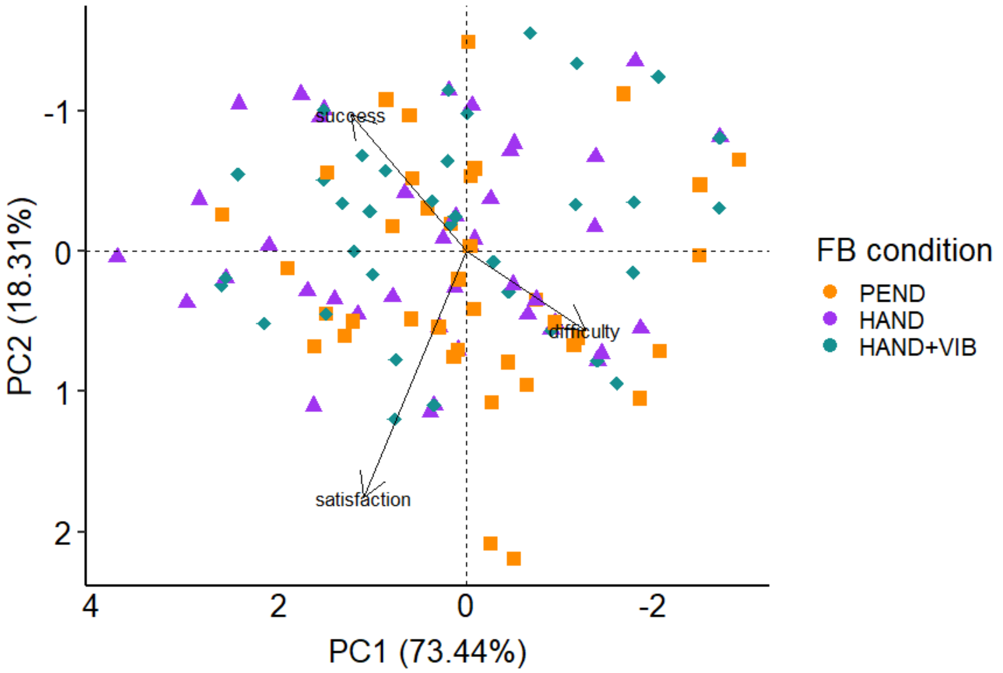


**Supplementary Figure 6.** PCA variables and individuals plot (right), PC1 is the selected user_experience score. FB conditions are color and shape-coded with orange squares, purple triangles and blue circles respectively representing PENDUL, HAND and HAND+VIB conditions.

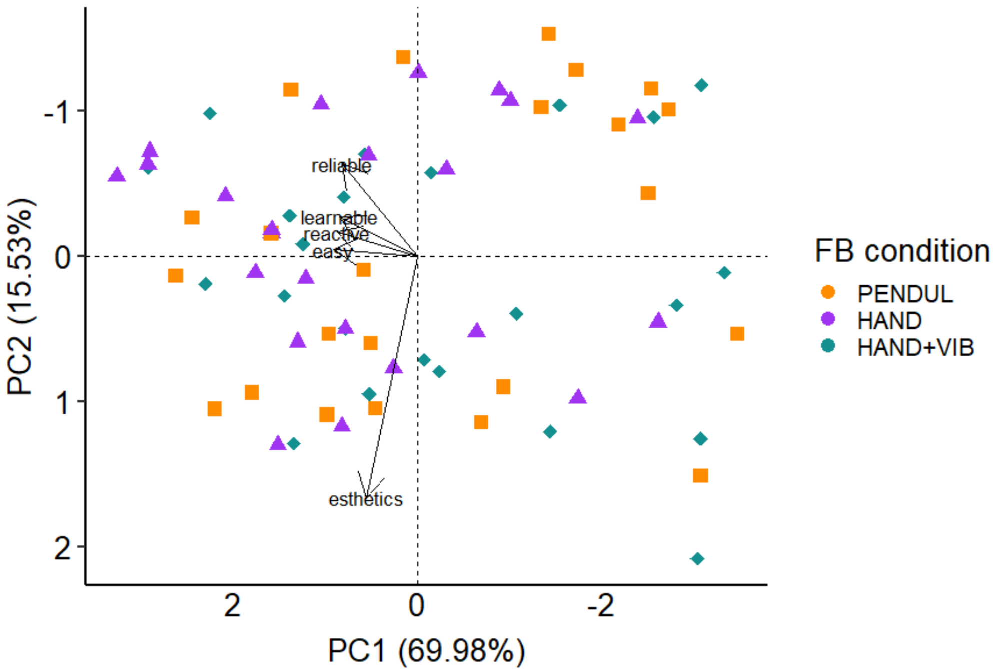


**Supplementary Figure 7**. PCA variables and individuals plot, PC1 is the selected feedback_usability score. FB conditions are color and shape-coded with orange squares, purple triangles and blue circles respectively representing PENDUL, HAND and HAND+VIB conditions.
